# Causal and directional elements of global brain dynamics

**DOI:** 10.64898/2026.05.27.726039

**Authors:** John Kochalka, Anish Mitra, Alexander D. White, Yu Wang, Kaiwen Sheng, Yukun Alex Hao, Chandan Kadur, Charu Ramakrishnan, Melissa Hernandez, Wendy Wenderski, Ki Eun Pyo, Nadya Andini, Thomas R. Clandinin, Michael Z. Lin, Shaul Druckmann, Karl Deisseroth

**Author notes:** Equal contribution (John Kochalka, Anish Mitra, Alexander D. White, Yu Wang).

## Abstract

Mammalian cognition appears to involve coordinated neural activity within and across brain-spanning networks. However, the governing principles of large-scale integration remain largely unknown. Here we developed an approach to record spontaneous cortex-wide neural activity (including with novel genetically encoded activity sensors), over very long timescales, and applied unbiased computation to capture conserved spatiotemporal regularities in these data. Initial screening extracted a set of directional elements: consistent spatiotemporal patterns of cortical activity that generalized across excitatory and inhibitory neuronal cell types as revealed by genetic targeting of activity sensors. In particular, novel genetically encoded voltage-sensing strategies revealed that the directional elements were not only highly conserved, but also were represented across the full neural activity frequency spectrum including the fastest timescales (gamma rhythms) enabled by voltage-sensing. Exploiting the spatiotemporal structure revealed by these directional elements, we tested for causal rules governing cortical network activation, using patterned optogenetic stimulation combined with activity imaging. We found that the directional propagation structure of these elements encoded a causal control hierarchy, as source regions (but not sink regions) sufficed to drive full element recruitment—a principle that held across all tested directional elements. Employing a panel of psychotropic drugs, we showed that directional element structure and excitability were robust to manipulations of neural and behavioral state, even as spontaneous network dynamics were reshaped in compound-specific ways. Finally, we developed and applied an all-optical sensing/control approach targeting the directional elements in a behavioral visual detection paradigm, revealing contributions of these conserved large-scale dynamics to elevated sensorimotor behavioral performance.

## Introduction

Orderly and sequential activation of neural circuits appears to be a fundamental principle of brain organization, across spatial and temporal scales. For example, at the scale of individual neurons, hippocampal place cell sequences active during behavior are replayed and preplayed during rest^1–4^, while choice-specific sequences have been identified in neocortical populations^5^. At the scale of cortical columns and local networks, spontaneous activity unfolds as structured ensemble sequences with repertoires closely mirroring those evoked by sensory stimulation^6–8^. At the scale of cortical areas, propagating waves of spontaneous activity gate perceptual behavior^9^ and recapitulate the spatiotemporal patterns elicited by peripheral stimulation^10,11^, and recent work has demonstrated that low-dimensional spatiotemporal structure underlies cortex-wide neural dynamics^12^. Across all these scales, the brain appears organized into functional elements with sequential co-activation, suggesting relevance of spatiotemporally-organized activity sequences in neural computation.

A parallel line of evidence establishes that the brain’s spontaneous activity architecture closely mirrors its task-evoked functional organization. For example, functional connectivity (measured hemodynamically with BOLD fMRI) in non-task-related resting states predicts aspects of task-evoked network recruitment as well as individual differences in cognitive performance across a range of paradigms^13–15^. Leveraging the correspondence between spontaneous and task activations, resting state fMRI has been used to identify a set of distributed functional networks involved in vision, audition, somatomotor function, attention, language, executive control, vigilance, memory, and self-referential cognition^16,17^. These networks are highly reproducible^18^ and identifiable in individual human subjects^19^. Further, similar network organization has been observed in non-human primates^20^ and rodents^21^, suggesting generality of these aspects of large-scale functional organization of the mammalian brain. However, by relying predominantly on zero-lag spatial correlations (rather than correlations across time), this literature has largely collapsed the temporal dimension of network dynamics—treating networks as spatial maps rather than spatiotemporal sequences– while the temporal order with which regions in a network are recruited may be important for understanding causal relationships across the brain, and for discovering how brain state shapes behavior in real time. Moreover, limitations of these established methods have made it challenging to establish how network activation patterns may relate to modern measures of cell type diversity, anatomical connectivity, the full spectrum of fast neural dynamics, and casual control hierarchies (beyond correlational signatures) relevant to brainwide dynamics and behavior.

Thus, despite pioneering advances over many years, fundamental questions about large-scale brain activity organization remain unanswered. Here we addressed these questions, carrying out an extensive unbiased screen of brain activity with novel optical approaches, combining long timescale recording in mice over many tens of hours in a manner that also spanned dorsal cortex in truly simultaneous fashion without scanning or tiling. This unbiased optical/computational screen revealed a set of robust and conserved directional elements: spatiotemporal patterns of cortical activation that generalized across excitatory and inhibitory cell types, across the full spectrum of cortical dynamics from delta to gamma bands, and across a broad range of pharmacological perturbations. Optogenetic testing revealed that the directional propagation structure of these elements was causally and behaviorally relevant, as activating source regions sufficed to drive network-wide recruitment while activating sink regions did not, and directional element activation in the moments preceding a visual stimulus causally determined whether that stimulus could be behaviorally detected. Together, the set of directional elements emerging from unbiased optical/computational screening establish a detailed and direct link between spontaneous local dynamics, brainwide activity in distributed cortical circuits, and moment-to-moment sensorimotor performance.

## Results

### Cortex-wide spontaneous activity screen reveals conserved directional elements

To study the large-scale organization of mammalian brain activity we initially performed widefield Ca^2+^ imaging across the entire dorsal cortices of head-fixed, awake mice using a custom tandem-lens macroscope (**Fig. 1a-b**)^22^. We began with Cux2-CreER;Ai148 transgenic mice^23,24^ expressing GCaMP6f in superficial excitatory neurons recording 12.6 h of spontaneous activity across 63 sessions in three animals. Cortical activity measured in this way exhibited large ongoing fluctuations in the absence of external sensory drive **(Fig. 1c, Supplementary Video 1)**. Seed-based correlation analysis revealed coupling not only locally and homotopically across hemispheres, but also to functionally related cortices– for example from vibrissal primary somatosensory cortex (vS1) to vibrissal motor cortex (vM1; **Fig. 1d**). Temporal cross-correlation analysis further revealed that vS1 activity consistently preceded vM1 by approximately 30 ms **(Fig. 1e)**, motivating the need for an approach that could capture both spatial and temporal structures of network activation.

**Figure 1.**
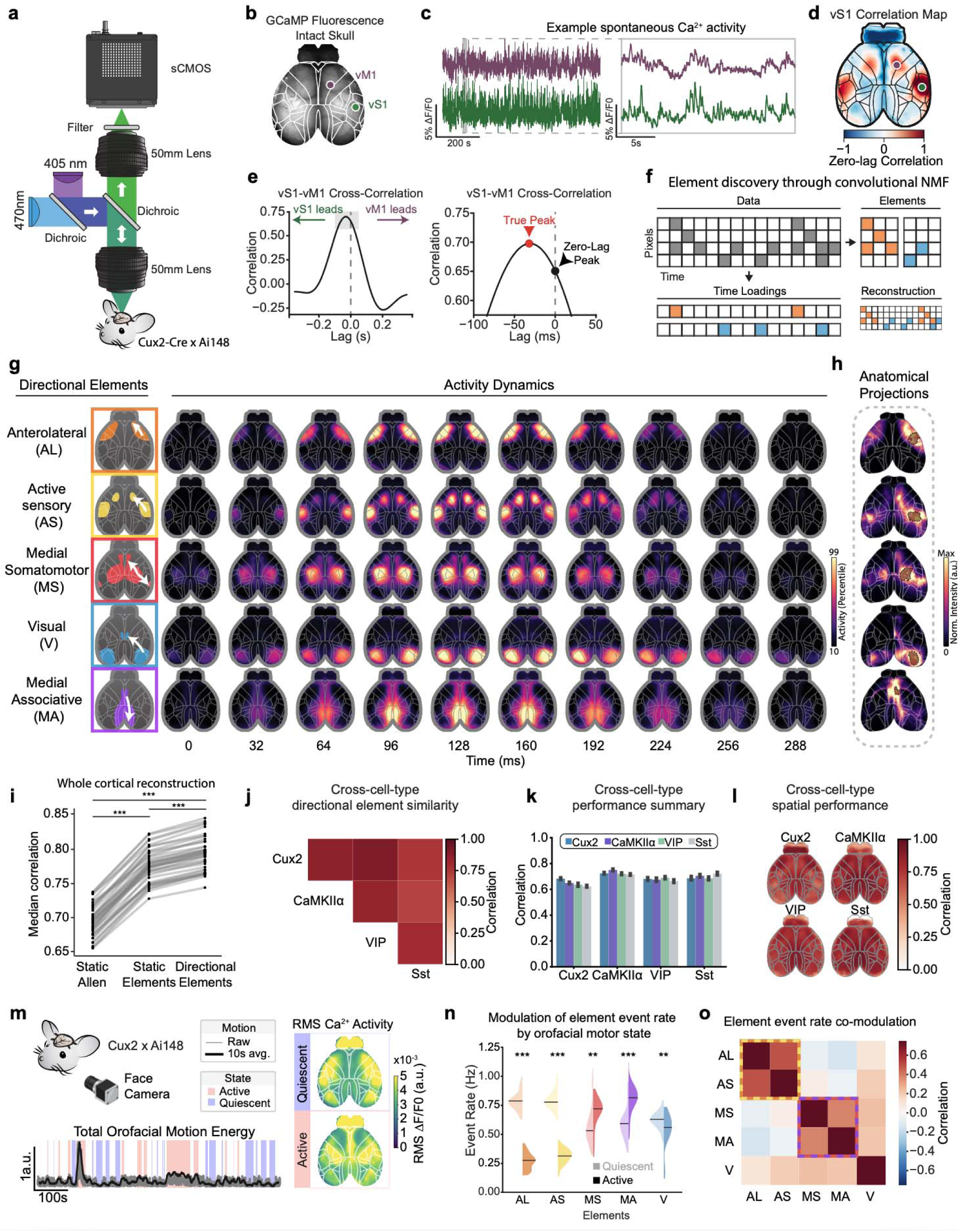
Cortex-wide spontaneous activity screen reveals set of spatiotemporal (directional) elements conserved across neuronal layers and cell types. **a.** Tandem lens widefield macroscope for recording Ca^2+^ signals from mouse dorsal cortex. **b.** Cleared-skull preparation enabling dorsal cortex-wide imaging of activity through the intact skull. **c.** Processed fluorescence traces from motor (vM1) and sensory (vS1) ROIs over a single 12 min recording of spontaneous activity (*left*), zoomed into the thirty-second grey highlighted area (*right*). **d.** Cortex-wide correlation map computed from a seed in vS1 highlighting a distributed, multi-area network encompassing vS1 and vM1. **e.** Cross-correlation between vS1 and vM1 timeseries from (c) (*left*), zoomed in to demonstrate the ∼30ms lag between vS1 and vM1 activity. **f.** Convolutional non-negative matrix factorization of Ca^2+^ data. Frames were vectorized into a two-dimensional pixel × time data matrix; seqNMF was then used to discover the consistent spatiotemporal (directional) elements and their respective temporal loadings. **g.** Directional elements derived from consensus clustering of n=63 recording sessions from N=3 Cux2-CreERT2 × Ai148 mice. **h.** Analysis of Allen Institute connectivity data^26,27^. Long-range connectivity profiles from directional element nodes exhibited correspondences to the identified directional elements. **i.** Cortex-wide activity reconstruction accuracy increased stepwise across three models: the Allen CCF parcellation, static elements derived from the temporal maximum projection of the discovered directional elements (i.e. their peak spatial pattern, discarding temporal dynamics), and the full dynamic directional elements. Each step yielded significantly improved reconstruction performance (N=3 mice, n=63 recordings, paired t-test, *** p < 0.001). **j.** Comparison of directional element similarity across cell types. Cross-correlation analysis of cell-type-specific directional elements revealed strong correspondence among all matched pairs. (One-sample t-test against zero, n=6 elements per cell-type group pair, all *p* < 0.001, FDR-corrected). **k.** Dataset-wise summaries of cross-cell-type directional element performance. Datasets from each cell type were fit with matched and non-matched cell-type specific directional elements. Distributions of pixelwise median correlation coefficients across datasets demonstrate each set of cell-type-specific networks performs well in reconstructing data from other cell types. (Comparison of matched and average of non-matched fits with two one-sided t-tests [TOST] procedure with equivalence bounds of ±0.10% of the matched fit; Cux2 n=63, CaMKIIα n=59, VIP n=70, Sst n=76, all *p* < 0.001). **l.** Spatial summaries of cross-cell-type directional element performance. Pixelwise maps of the groupwise median correlation coefficient show strong, spatially consistent ability of non-matched elements to model data from each cell type. (Pixel-wise comparison of matched and average of non-matched fits with TOST procedure with equivalence bounds of ±0.10% of the matched fit to determine fraction of matched pixels: Cux2: 84.4%, CaMKIIα: 90.1%, VIP: 94.8%, Sst: 86.4% of pixels; Binomial test on the fraction of matched pixels, all *p* < 0.001; Cux2 n=63, CaMKIIα n=59, VIP n=70, Sst n=76) **m.** Analysis of spontaneous behavior. Total orofacial motion energy was computed for each frame; a 10-second moving average was then used to delineate Active and Quiescent periods. Maps of the Root-Mean-Square (RMS) ΔF/F_0_ activity revealed state-dependent differences in cortical activity. (Pixel-wise paired t-test on Active vs Quiescent RMS followed by FDR correction. See **Extended Data Fig. 4f**). **n.** Directional element event rates as a function of behavioral state. AL/AS rates were strongly reduced during the Active state (Cohen’s *d’s* AL: -4.1, AS: -3.75), V rates were moderately suppressed (Cohen’s *d* -0.45), and MS/MA exhibited increased activity (Cohen’s *d’s* MS: 0.41, MA: 0.92). Paired *t-*test per directional element (** *p* < 0.01, *** *p* < 0.001). **o.** Directional element event rate correlation matrix showing block structure, with AL/AS forming one cluster and MS/MA forming a second.

To identify the dominant spatiotemporal patterns of cortical dynamics, we applied seqNMF^12,25^, a convolutional factorization algorithm that decomposes activity movies into a set of spatiotemporal factors (“elements”) and their temporal loadings **(Fig. 1f)**. Applying a group consensus element discovery pipeline **(Extended Data Fig. 1)** to find the dominant, shared spatiotemporal patterns of cortical activity in the Cux2 dataset yielded a small set of directional elements **(Fig. 1g)**: an Anterolateral (AL) element initiating in orofacial somatosensory areas and propagating anteromedially through primary and secondary motor cortices; an Active Sensing (AS) element originating in vS1 and recruiting vM1; a Somatomotor (SM) element spreading from hindlimb, forelimb, and trunk areas to secondary motor (M2) and somatosensory areas (S2); a Visual (V) element sweeping through primary (V1) and higher order visual (HOV) cortices to dorsal anterior cingulate cortex (dAC); a Medial Associative (MA) element propagating from medial prefrontal (mPFC) to retrosplenial cortex; and an Olfactory (OB) element dominated by quasi-periodic olfactory bulb activation. We focus here primarily on the first five directional elements of the six emerging from the unbiased algorithm, deferring detailed analysis of OB to the Supplemental Information as OB was confined largely to the olfactory bulb without major propagation to neocortex.

To explore the relationship between the discovered directional elements and underlying structural connectivity, we analyzed data from the high-quality Allen Institute Mouse Brain Connectivity Atlas^26,27^. Selecting experiments with injection sites near the source regions of each cortical network (**Extended Data Fig. 2**; see Methods), we averaged the maximum-projection fluorescence signal across experiments within each network and compared the resulting projection patterns to the spatial footprints of the corresponding network elements. Injections at source regions produced anterograde labeling that closely mirrored each network element, with projection targets aligning to regions that activated later in the spatiotemporal sequence **(Fig. 1h)**. Notably, injections at sink regions revealed a reciprocal pattern **(Extended Data Fig. 2)**, suggesting that the directed nature of cortical dynamics — stereotyped patterns where source activity leads the rest of the network — cannot be fully explained by projectomic anatomical connectivity and requires activity recording.

To quantify the representational advantage conferred by capturing cortical dynamics, we compared three levels of parcellation complexity in their ability to reconstruct Cux2 imaging data: standard Allen CCFv3 anatomical parcels, "static" elements derived by taking the maximum projection of each directional element across time, and the fully dynamic directional elements themselves **(Fig. 1i)**. Each successive level provided a significant improvement in data fit: static elements outperformed Allen parcels (p < 0.001), and dynamic directional elements outperformed static elements (p < 0.001). This hierarchy demonstrates that functional definitions of cortical regions— even when restricted to zero-lag spatial footprints that ignore propagation— outperform cytoarchitectonic boundaries in capturing spontaneous cortical variance; crucially, restoring the full temporal propagation structure of directional elements yields a further significant gain, indicating that a substantial fraction of spontaneous activity is organized along dynamic, directionally-specific axes that are invisible to both anatomy and static functional parcellations.

To test whether these directional elements were a specific feature of superficial excitatory neurons or a general property of cortical circuits, we repeated the spontaneous imaging and element discovery protocol in separate cohorts of mice expressing GCaMP in vasoactive intestinal peptide (VIP)-expressing interneurons (N=3, 14 h), somatostatin (Sst)-expressing interneurons (N=3, 15.4 h), and CaMKIIα-expressing excitatory neurons spanning LII/III and L5 (N=6; 11.8 h)^24^. In all cell-type cohorts, the same six directional elements were recovered with strikingly similar spatiotemporal structure **(Extended Data Fig. 3; Supplementary Video 2)**. Pairwise cross-correlation peaks between matched elements across cell types averaged near unity **(Fig. 1j)**. To further assess generalization, each dataset was fit with all four sets of cell-type-specific directional elements; in every case, the non-matched elements fit the data comparably to the matched set (**Fig. 1k**, all non-matched fits within ±10% of matched fits, two one-sided *t*-tests [TOST] *p* < 0.001), indicating that any of the four groups of directional elements captures the dominant structure of spontaneous activity regardless of the recorded cell type. To assess spatial generalization, we computed pixel-wise correlation for all cross-cell-type fits; across all four cell-type groups, the large majority of brain pixels showed non-matched fits within ±10% of the matched fit (Cux2: 84.4%, CaMKIIα: 90.1%, VIP: 94.8%, Sst: 86.4% of pixels; **Fig. 1l**). In each case, this proportion far exceeded the 5% false-positive rate expected by chance, as confirmed by a one-sided binomial test (*p* < 0.001 for all groups), indicating that non-matched directional elements produce fits that are spatially commensurate with matched elements across the overwhelming majority of dorsal cortex.

To characterize the relationship between spontaneous network dynamics and behavior, we recorded high-speed videography and quantified motor behavior via frame-wise motion energy^28,29^, segmenting epochs into high and low motor activity periods **(Fig. 1m)**. RMS ΔF/F_0_ maps revealed clear state-dependent reorganization of cortical activity, with quiescent periods characterized by elevated variance in V1, vS1, and anterolateral sensorimotor cortices, and active periods showing elevated variance in trunk, forelimb, hindlimb, and HOV regions. Directional event rates were similarly state-dependent: AL, AS, and, to a lesser extent, V, rates were suppressed during active periods, while MS and MA showed elevated rates **(Fig. 1n)**. Analysis of event rates in a separate cohort adapted to running on a wheel during spontaneous imaging recapitulated the collective organization revealed by analysis of bulk orofacial movement **(Extended Data Fig. 4)**. Pairwise correlations among event rates further revealed a block-diagonal structure — AL and AS co-varied strongly with one another, as did MS and MA, with weak to negative cross-group correlations **(Fig. 1o)**. This collective organization was recapitulated by analysis of faster-timescale, finer-grained behavioral features such as snout and whisker pad movements as well as pupil size and saccades **(Extended Data Fig. 4)**. Taken together, these results suggest the cortex tends to dwell in either an AL/AS-dominant or MS/MA-dominant regime, echoing the task-positive/task-negative anticorrelation described in resting-state fMRI^30^.

Together, these results establish a compact, conserved vocabulary of large-scale cortical dynamics in the mouse. Six spatiotemporal directional elements—spanning sensorimotor, visual, medial associative, and olfactory systems—reproducibly captured the dominant structure of spontaneous dorsal cortical activity, and did so with comparable fidelity whether recorded from superficial excitatory neurons, deep excitatory neurons, or either of two major inhibitory interneuron subtypes. This conservation across cell types is notable, since given the well-documented distinct physiological properties and computational roles of VIP^31^, Sst^32^, and pyramidal neuron populations^33^, one might have expected each to exhibit a unique large-scale organizational signature. Instead, these results suggest that at the level of dominant population-wide dynamics, cortical location rather than cell identity is the primary determinant of network membership. This motivates treating these six directional elements as a cell-type-agnostic reference atlas of mouse cortical network architecture—a foundation on which subsequent analyses of behavioral correlates and causal control can be built.

### Source but not sink regions suffice to recruit directional elements

The spatiotemporal structure of directional elements revealed consistent vectorial propagation of activity from "source" regions (early-active areas) to "sink" regions (late-active areas). We hypothesized that this structure reflects a fundamental asymmetry in cortical network control, whereby source activation should suffice to recruit the full network, while sink activation should not drive upstream responses. To test this hypothesis directly, we constructed a novel all-optical system enabling simultaneous patterned one-photon optogenetic stimulation and widefield Ca^2+^ imaging **(Fig. 2a)**. A digital micromirror device (DMD) coupled to a 577 nm laser enabled spatially precise cortical activation, validated by projection of arbitrary illumination patterns onto a test surface **(Fig. 2b)**. Mice co-expressed GCaMP8m and frChRmine—a red-shifted, low-conductance channelrhodopsin we had developed via high-resolution ChRmine structural work^34^, chosen for its minimal spectral overlap with GCaMP excitation wavelengths (here 470 nm) and fast kinetics **(Fig. 2d)**, all under the CaMKIIα promoter via AAV-PHP.eB retro-orbital injection^35–37^. Single-trial data confirmed that a brief optical pulse (6 ms, 0.6 mm² ROI) reliably recruited the targeted cortical site despite substantial variability in ongoing activity across trials, with robust responses evident on individual trials as well as in the 25-trial average **(Fig. 2c)**.

**Figure 2.**
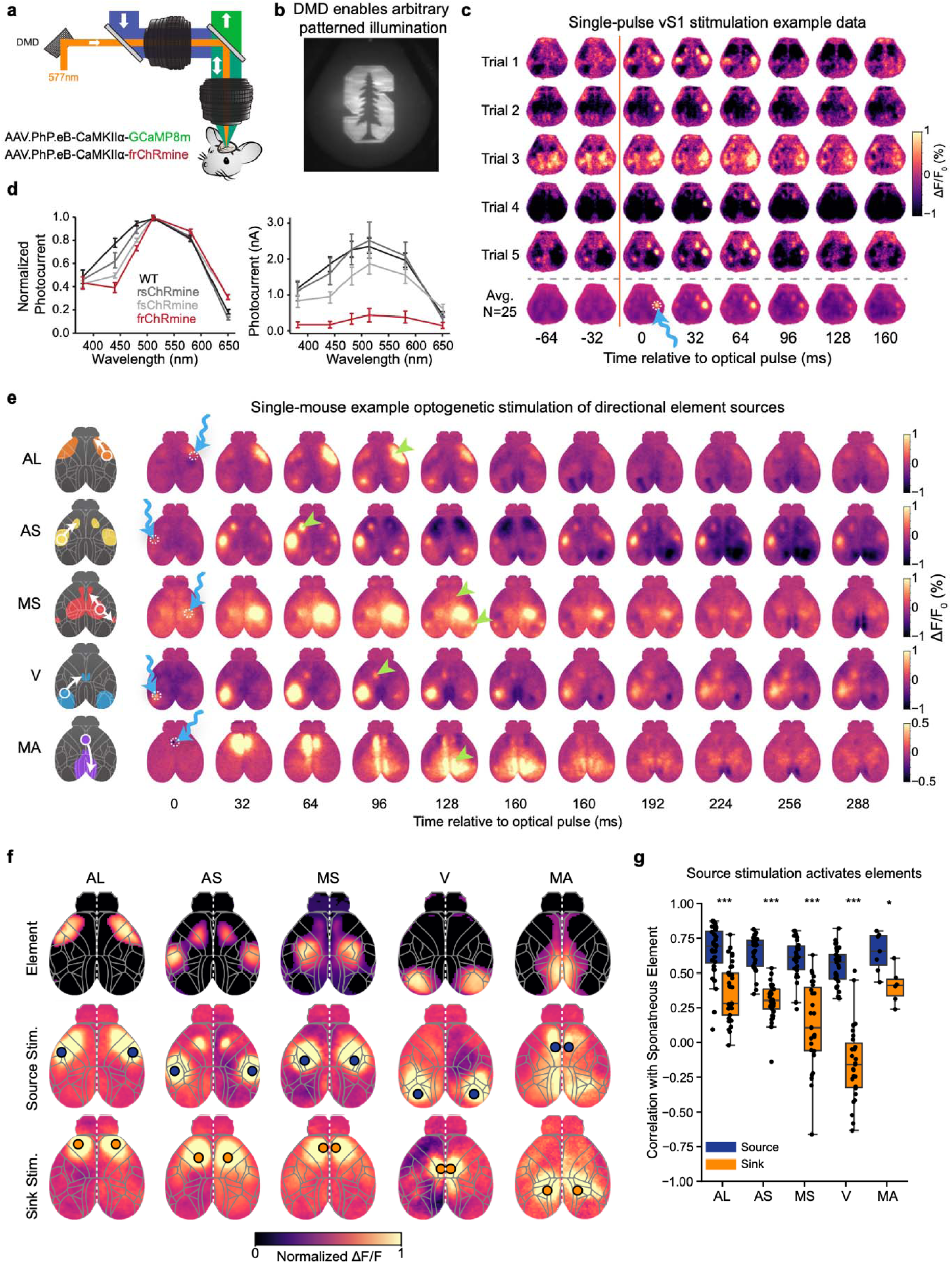
Optogenetic stimulation of source areas suffices to recruit full cortex-spanning directional elements. **a.** Schematic of microscope enabling all-optical assays using patterned one-photon illumination in a red-shifted channel to modulate neural activity during simultaneous Ca^2+^ imaging. Brain-wide expression of GCaMP8m and frChRmine in CaMKIIα neurons was achieved through retro-orbital injection of AAV-PhP.eB in wild-type C57BL/6 mice. **b.** Widefield image of tree logo demonstrating spatial patterning capabilities of the system (912 × 1140 micromirrors with 7.6 µm pitch, Active area: 6.1614 mm × 9.855 mm, magnification 1:1 to sample). **c.** Single-trial and trial-averaged (n=25) evoked responses following patterned optical stimulation of a site located in vS1, displayed in the native imaging space from a single technical replicate. The vertical orange bar denotes timing of delivery of a 6 ms optical pulse to a 0.6 mm^2^ ROI denoted by the wavy blue arrow (*bottom row*). We observed robust recruitment of the targeted ROI despite variability in ongoing activity (e.g. compare Trial 3 vs Trial 2), with frequent but not deterministic recruitment of homotopic vS1 and downstream vM1. **d.** Normalized and absolute photocurrent action spectra of ChRmine variants highlighting the more red-shifted yet comparatively weak responses of frChRmine in contrast to other variants. **e.** Single replicate example data for directional element source stimulation. Each row is a series of baseline-corrected ΔF/F_0_ images averaged over 25 trials in an individual mouse. Blue wavy arrows indicate target of optogenetic stimulation. Green arrows denote downstream sites of recruited activity. **f.** Spatial maps of patterns evoked by focal optogenetic stimulation. *Upper*: maximum projections of each directional element, for visual comparison. *Middle*: average evoked response from optogenetic stimulation of the source of each respective directional element (AL n=32 N=15; AS n=33 N=16; MS n=27 N=10; MA n=7 N=2; V n=33 N=16). *Lower*: as above but for the directional element sinks (AL n=33 N=16; AS n=33 N=16; MS n=27 N=10; MA n=6 N=6; V n=27 N=10). **g.** Analysis of similarity between optogenetic-stimulation-evoked responses and directional elements. The first four frames of the trial-averaged response were averaged, warped to Allen CCFv3 space, then spatially correlated with the maximum projection of the corresponding directional element, restricting the analysis to the stimulated hemisphere. Boxplots show the median, IQR, and whiskers indicating 1.5 × IQR. (Two-way ANOVA, main effect of source vs sink *p* < 0.001, post-hoc *t*-test of source vs sink per element, * = *p* < 0.05, ** = *p* < 0.01, *** = *p* < 0.001).

We defined source and sink ROIs for each of the five cortical network elements based on the early- and late-active regions of the corresponding spontaneous activity pattern (**Extended Data Fig. 2**; Methods), and stimulated each site while imaging cortex-wide responses. Single trials revealed that source stimulation of the Anterolateral (AL), Active Sensory (AS), Medial Somatomotor (MS), Visual (V), and Medial Associative (MA) network elements each elicited a spatiotemporal pattern of downstream cortical recruitment that closely recapitulated the structure predicted by the corresponding spontaneous activity pattern **(Fig. 2e)**. Summary maps confirmed this pattern at the group level across animals: source stimulation produced activity that propagated through the expected network territory, while sink stimulation showed no evidence of upstream recruitment **(Fig. 2f, Supplementary Video 3)**. Quantification of the spatial correlation between optogenetically evoked patterns and the corresponding spontaneous network element **(Extended Data Fig. 5)** confirmed that source stimulation produced significantly higher similarity than sink stimulation across all five networks tested (**Fig. 2g**; two-way ANOVA, main effects of network and source vs sink both p < 0.001; post-hoc *t*-tests: AL *p* < 0.001, AS *p* < 0.001, MS *p* < 0.001, V *p* < 0.001, MA *p* < 0.05, Bonferroni corrected), establishing causal underpinnings of directional element spatiotemporal structure.

### Psychotropic modulation of directional element utilization with preserved topology

Having identified spatiotemporal structure and control principles of cortical directional elements under baseline conditions, we next asked how robust these dynamical features were to perturbations of brain and behavioral state. We began with three psychotropic drugs characterized by distinct mechanisms of action: ketamine (50 mg/kg; known actions primarily via excitatory and neuromodulatory receptor action), diazepam (2 mg/kg; known actions primarily via inhibitory receptor action), and amphetamine (2 mg/kg; known actions primarily via neuromodulatory transporter action).

Alongside saline controls, we performed widefield Ca² imaging continuously before, during, and after drug administration. Examination of RMS ΔF/F spanning 5–35 minutes post-injection) revealed drug-specific reorganization of bulk cortical activity, with each compound producing a distinct spatial signature **(Fig. 3a)**. However, despite these differences in overall activity patterns, directional elements discovered under each drug condition were remarkably similar to their saline counterparts **(Extended Data Fig 6, Supplementary Video 4)**. Cross-correlation analysis revealed high drug-vs-saline similarity (>∼80% correlation) for each directional element, for all three compounds, with the notable exception of reduced cross-correlation of the MA element under ketamine **(Fig. 3b).** Examining spatiotemporal dynamics of the ketamine-specific MA recruitment pattern revealed qualitatively distinct dynamics in retrosplenial cortex, including a novel initiation site in lateral agranular RSP and global waves following prominent RSP activation—features absent in the saline condition **(Fig. 3c)**.

**Figure 3.**
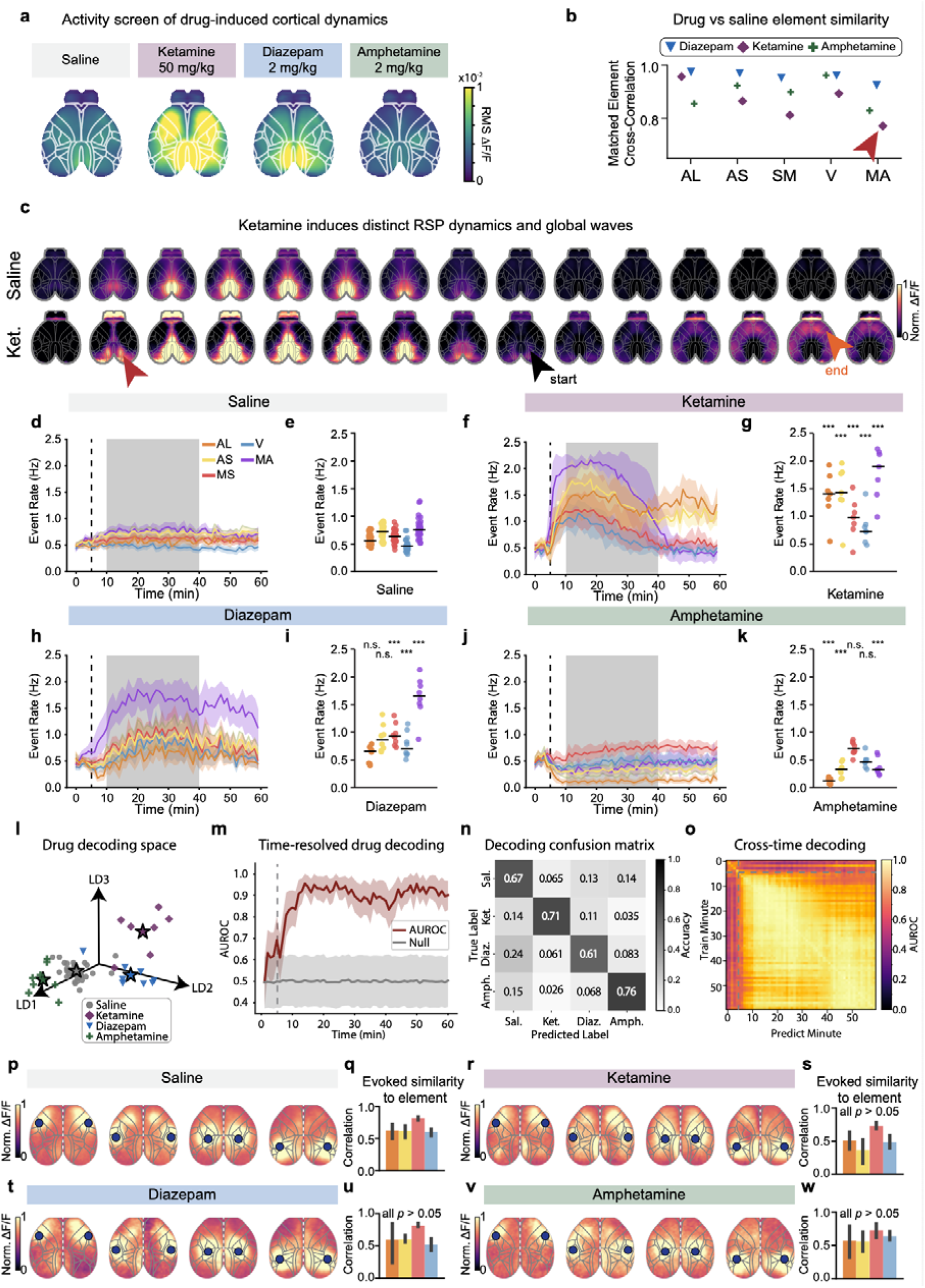
Pharmacological states modulate element utilization while preserving topology and excitability. **a.** Brain-wide activity screen following administration of ketamine, diazepam, and amphetamine at doses indicated. RMS ΔF/F_0_ is mapped covering the peak drug phase (5-35 mins following administration) revealing coarse-grained systems-level differences in drug-induced neural activity. **b.** Directional elements derived from peak drug phases were highly similar; cross correlation comparing matched pairs of drug-specific basis motifs to the saline-derived set reveals strong correspondence, with the red arrow highlighting the largest difference seen (altered Medial Associative or MA dynamics under ketamine). **c.** MA dynamics are altered under Ketamine. First red arrow highlights distinctive initiation site of ketamine-induced RSC oscillation in lateral agranular RSC. Black arrow shows start of spreading wave dynamics emanating from RSC anterolaterally across cortex following MA event, while orange arrow denotes the end of the wave. **d-k,** Time resolved directional element event rates following drug injection at 5 mins (gray dashed line). Time courses of directional element event rates before and after administration of (d) saline (n=33, N=18), (f) ketamine (N=7), (h) diazepam (N=8), or (j) amphetamine (N=8). Shaded region indicates the peak drug phase (5–35 min post-administration). Traces are plotted as mean ± 95% confidence interval. Summary panels show statistics of directional element event rates during the peak drug phase for saline (e), ketamine (g), diazepam (i), and amphetamine (k). Post hoc pairwise comparisons (Tukey HSD) were performed following a one-way ANOVA across drug conditions for each directional element. Only comparisons versus saline are shown; asterisks indicate significance levels: * *p* < 0.05, ** *p* < 0.01, *** *p* < 0.001. **l–o)** Decoding drug identity from directional element dynamics. **l,** Example Linear Discriminant Analysis (LDA) subspace showing projections of dataset-level summaries onto the first three discriminant components, colored by drug condition, revealing clustering with apparent linear separability between conditions. **m,** Time-resolved LDA decoding performance (Area Under the Receiver Operating Characteristic Curve, AUROC) demonstrating robust and sustained discrimination of drug identity following administration. Traces are plotted as mean ± 95% confidence interval. **n,** Confusion matrix summarizing classification performance across drug conditions. Each cell represents the fraction of trials in which the drug indicated by the row was classified as the column label. The classifier performed above chance for all four conditions, with highest accuracy for amphetamine (0.76) and ketamine (0.71), followed by saline (0.67) and diazepam (0.61). The most prominent off-diagonal error was diazepam being misclassified as saline (0.24); all other off-diagonal values were small (≤0.15), indicating that the remaining drug conditions were largely distinguishable from one another. **o,** Cross-time decoding analysis, in which classifiers trained at a given time point (rows) are tested across all other time points (columns). Color indicates AUROC, with bright yellow reflecting near-perfect classification and dark purple reflecting chance-level performance. The broadly high AUROC values across the majority of the matrix (outside the first ∼5 minutes pre-injection, dashed lines) indicate that drug-specific network signatures are temporally stable— a classifier trained at nearly any time point generalizes well to nearly any other, suggesting that the neural features distinguishing drug conditions are sustained rather than transient. **p–w)** Spatial patterns of activity evoked by focal optogenetic stimulation of directional element-defined source sites across pharmacological conditions. Maps of stimulation-evoked activity shown following activation of source nodes during (p) saline (n=20, N=7), (r) ketamine (n=6, N=3), (t) diazepam (n=4, N=2), and (v) amphetamine (n=6, N=3) conditions. Panels q,s,u,w show summary quantification of similarity between evoked activity patterns and the corresponding endogenous directional elements for saline (q), ketamine (s), diazepam (u), and amphetamine (w). Error bars represent ± 95% confidence interval. Two-way ANOVA for effects of drug (*p* = 0.66), element (*p* < 0.001) and their interaction (*p* = 0.014) followed by post-hoc pairwise *t*-tests versus saline with Holm correction within each element and Bonferroni correction across elements (all *p* > 0.05 after correction).

Analysis of directional element event rates over time **(Extended Data Fig 7)** revealed distinct signatures for each drug **(Fig. 3d–k)**. Saline injection produced no systematic change in event rates, which remained stable and evenly distributed across all directional elements throughout the session **(Fig. 3d–e)**. Ketamine produced a dramatic and sustained elevation of event rates across all directional elements beginning shortly after injection, with a particularly prominent increase in MA that persisted throughout the administration and post-administration phases (**Fig. 3f–g**; all elements *p* < 0.001 vs. saline, Cohen’s *d*’s AL: 4.11, AS: 3.15, MS: 1.90, V: 2.08, MA: 4.10). Diazepam produced a more selective elevation, predominantly increasing MA event rates while leaving other elements comparatively less affected (**Fig. 3h–i**; MS, V, MA *p* < 0.001, others n.s.; Cohen’s *d*’s AL: 0.28, AS: 1.16, MS: 2.51, V: 2.24, MA: 3.78). In contrast, amphetamine exhibited a suppressive effect, markedly reducing AL, AS, and MA event rates while leaving MS and V directional elements relatively intact (**Fig. 3j–k**; AL, AS, MA *p* < 0.001, others n.s.; Cohen’s *d*’s AL: -5.15, AS: -3.39, MS: 0.74, V: 0.01, MA: -2.36).

These drug-specific signatures in directional element event rates suggested that the multi-dimensional state of cortical dynamics might carry sufficient information to identify the drug condition from directional element activity alone. To test this idea, we trained Linear Discriminant Analysis (LDA) decoders on each minute of time-resolved event rate data. Visualization of the LDA subspace revealed clear separation of the four conditions **(Fig. 3l, Extended Data Fig. 7)**. Time-resolved decoding showed that classification accuracy rose rapidly following injection, reaching high performance (AUROC >0.9) within approximately 10 minutes and remaining stable throughout the session **(Fig. 3m)**. A confusion matrix revealed accurate decoding across all four conditions, with no systematic pattern of confusability between drugs **(Fig. 3n)**. A cross-time decoding analysis revealed stability of the decoding subspace from shortly after drug injection onward, with high cross-time generalization persisting across the remainder of the recording **(Fig. 3o)**.

Given altered directional element incidence rates, we then asked whether these potent drug-mediated perturbations of spontaneous network dynamics might reflect altered element “excitability”, in the sense of altered susceptibility to causal elicitation of directional elements via source stimulation. We repeated our all-optical assay during the peak drug phase for saline, ketamine, diazepam, and amphetamine, examining optogenetically evoked responses from AL, AS, MS, and V source sites. Across all four drug conditions, source stimulation continued to recruit expected downstream network activity, with robust spatial similarity to the spontaneous directional element that was statistically indistinguishable from the saline condition (**Fig. 3p–w**; no significant differences in evoked similarity scores across drug conditions). These results revealed indicate that while psychotropic drugs profoundly reshape the statistics of spontaneous network activity—altering which directional elements are most active and at what rates—these highly potent and clinically-relevant agents leave intact the capacity of underlying circuit architecture at source regions to drive downstream element recruitment. Thus we have identified a dissociation between two aspects of cortical network organization: the statistics of spontaneous directional element incidence are highly sensitive to pharmacological perturbation, while underlying architecture within directional elements is remarkably resilient. Ketamine, diazepam, and amphetamine each produced distinct and dramatic reorganizations of spontaneous network element event rates—signatures so reliable that the drug condition could be decoded from cortical dynamics alone with high accuracy within minutes of injection. Yet despite these profound differences in spontaneous activity, the ability of source regions to recruit downstream network activity was preserved across all conditions tested, suggesting that the source-to-sink control principle identified herein reflects a stable property of cortical structure rather than a contingent feature of particular dynamical states.

These findings carry two broader implications. First, the drug decoding results establish network element event rates as a sensitive and interpretable readout of global brain state—one that may prove useful as a translational bridge between preclinical pharmacology and human neuroimaging, where drug effects on resting-state network dynamics are increasingly used as biomarkers of target engagement^38^. Second, the robustness of optogenetically evoked network recruitment under pharmacological challenge raises the possibility that source-targeted stimulation could restore normal patterns of network activity even when spontaneous dynamics are severely disrupted—a potentially important consideration for therapeutic neuromodulation approaches targeting network dysfunction in psychiatric and neurological disease^39–41^.

### Sensorimotor behavior casually modulated by pre-task directional element activity

Having established spatiotemporal structure, behavioral correlates, and pharmacological modulation properties of cortical directional elements, we next explored specific optogenetic recruitment of directional elements to test for potential causal modulation of behavior. We trained water-restricted mice on a visual detection task requiring licking in response to a faint drifting Gabor patch to earn a water reward, with early licks during the inter-trial interval punished by an air puff **(Fig. 4a–b)**. Mice acquired the task robustly, showing a sharp rise in lick rate following stimulus onset that decayed over subsequent seconds **(Fig. 4c)**.

**Figure 4.**
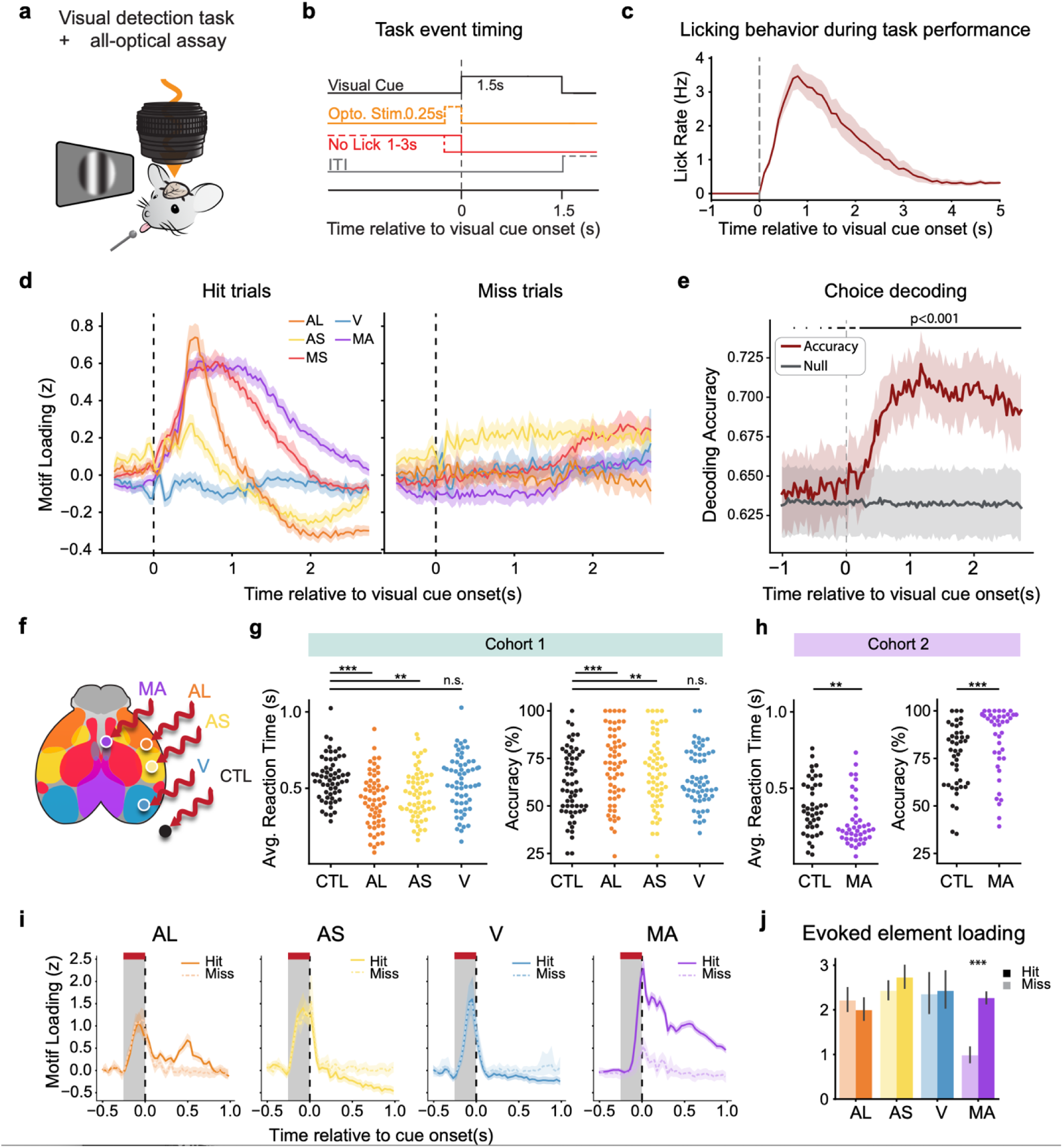
Sensorimotor behavior casually modulated by pre-task directional element recruitment. **a.** Visual detection task with simultaneous Ca^2+^ imaging and patterned optogenetic stimulation. Water-deprived mice were trained to detect a faint drifting Gabor patch against an isoluminant gray background and lick to receive a water reward. **b.** Task event structure. Following a variable intertrial interval (ITI), mice were required to withhold licking (“No lick”) for 1-3s before a visual cue presentation could be triggered. In a subset of trials, mice were optogenetically stimulated for 250 ms prior to visual cue onset, in which case the No Lick condition terminated at the onset of optogenetic stimulation (dashed line). **c.** Lick rate traces from non-optogenetic-stimulation trials show mice robustly increase licking in response to the visual cue. **d.** Visual-cue-triggered average z-scored temporal loadings for each directional element for Hit and Miss trials. Time courses show strong activation of MA, AL, and MS in Hit compared to Miss trials. Traces are plotted as mean ± 95% confidence interval. e. Choice decoding demonstrating robust separability of behavior-related brain states in the low-dimensional space of network elements. Traces are plotted as mean ± 95% confidence interval. **f.** Targets for optogenetic stimulation. Sources of the MA, AL, AS, and V directional elements, as well as an adjacent out-of-brain control site (CTL), were stimulated on subsets of trials immediately prior to stimulus onset. **g.** Effects of pre-stimulus directional element source stimulation on response latency. Trial-wise distributions of reaction time revealed significantly faster responses following AL, AS, and MA stimulation (Two-way ANOVA with factors mouse and trial type, main effect of mouse *p* < 0.01, main effect of trial type *p* < 0.05, interaction *p* > 0.05, followed by post-hoc *t*-test per trial type; * = *p* < 0.05, ** = *p* < 0.01, *** = *p* < 0.001). Effects of pre-stimulus directional element source stimulation on probability of stimulus detection. Significant performance improvements were observed following AL, AS, and MA stimulation (population-averaged logistic regression (GEE) with trial type as a fixed factor and repeated measurements clustered by mouse and session. Pre-stimulus AL, AS, and MA optogenetic stimulation yielded significantly higher odds of correct responses compared to Control. AL: *p* < 0.001, OR =1.42, 95% CI=1.17-1.72; AS: *p* < 0.01, OR=1.26, 95% CI=1.07-1.49, **h.** as in g but for MA vs control comparison (involving a different viral expression strategy so implemented in a separate Cohort 2 (Methods). MA: *p* < 0.001, OR = 1.97, 95% CI=1.51-2.57). **i.** Directional element temporal loadings on optogenetic stimulation trials as a function of trial outcome. All traces show little difference in the optogenetically-evoked network activity except MA. Traces are plotted as mean ± 95% confidence interval. **j.** Quantification of directional element recruitment on optogenetic stimulation trials for hits vs misses. Significance was observed only in MA (permutation test, *p* < 0.001).

We first characterized the relationship between directional element dynamics and behavioral outcome on non-optogenetically modulated trials. Peristimulus directional element loadings revealed different dynamics on hit versus miss trials: hit trials were characterized by robust directional element responses following visual cue onset, while miss trials did not, across all elements **(Fig. 4d)**. To quantify the discriminability of hit and miss neural states from the directional element basis alone, we trained a time-resolved LDA decoder on the element loading timeseries. Choice decoding performance rose significantly above chance prior to visual cue onset and climbed sharply following the cue, reaching approximately 70–75% accuracy in predicting behavioral response on held out hit or miss trials **(Fig. 4e)**, demonstrating that the directional element representation captures behaviorally relevant variance in cortical state. These results suggested the possibility that directional element recruitment could be involved in enhancing behavioral performance (although still requiring causal testing).

To causally test the contribution of specific directional elements to detection performance, we optogenetically stimulated the sources of the AL, AS, V, and MA elements in the 250 ms prior to visual cue onset on pseudorandomly interleaved trials, comparing with a control (out-of-brain) stimulation site **(Fig. 4f, Extended Data Fig. 8)**. Stimulation of AL, AS, and MA sources each produced significant improvements in task performance relative to control stimulation, reducing average reaction time (**Fig. 4g**; AL *p* < 0.001, AS *p* < 0.01, MA *p* < 0.01; V *p* > 0.05) and increasing detection accuracy (**Fig. 4h**; AL *p* < 0.001, AS *p* < 0.01, MA *p* < 0.001; V *p* > 0.05). In contrast, visual cortex source stimulation had no significant effect on either measure, indicating that the behavioral improvements were not simply a consequence of broadly increasing cortical excitability or providing an additional visual percept. Analysis of directional element recruitment on hit versus miss trials within each stimulation condition revealed a specific dependence on MA. Whereas other networks showed no relationship between the degree of optogenetic element recruitment and successful visual detection, on trials where optogenetic stimulation successfully recruited stronger MA directional element activity, mice were more likely to correctly detect the stimulus, while miss trials were associated with weaker MA recruitment **(Fig. 4i–j)**. This relationship suggested that the extent of MA directional element engagement in the pre-stimulus and peri-stimulus period plays a specific role in gating perceptual task performance.

### Voltage imaging reveals shared directional element structure across frequency bands

Genetically encoded Ca² sensors, as used here thus far, provide excellent signal-to-noise and cell-type specificity but are fundamentally limited in resolving fast neural dynamics due to slow Ca² extrusion ^42,43^. Voltage indicators report membrane potential fluctuations directly, enabling access to the full frequency content of neural activity from low-frequency delta fluctuations through fast gamma-band dynamics^44^. However, for questions such as those addressed here, voltage imaging approaches have been by limited by the low signal-to-noise ratios and rapid photobleaching of existing indicators^45–49^. To ask whether the directional network element organization we identified extends to the important and broad higher-frequency bands of neural activity, we repeated the cortex-wide spontaneous activity screen using a novel genetically encoded voltage indicator (GEVI), ASAP7y^50^, with absorption and emission spectra shown in **Fig. 5a**, imaged with a novel widefield macroscope offering substantially improved photon collection efficiency^51^.

**Figure 5.**
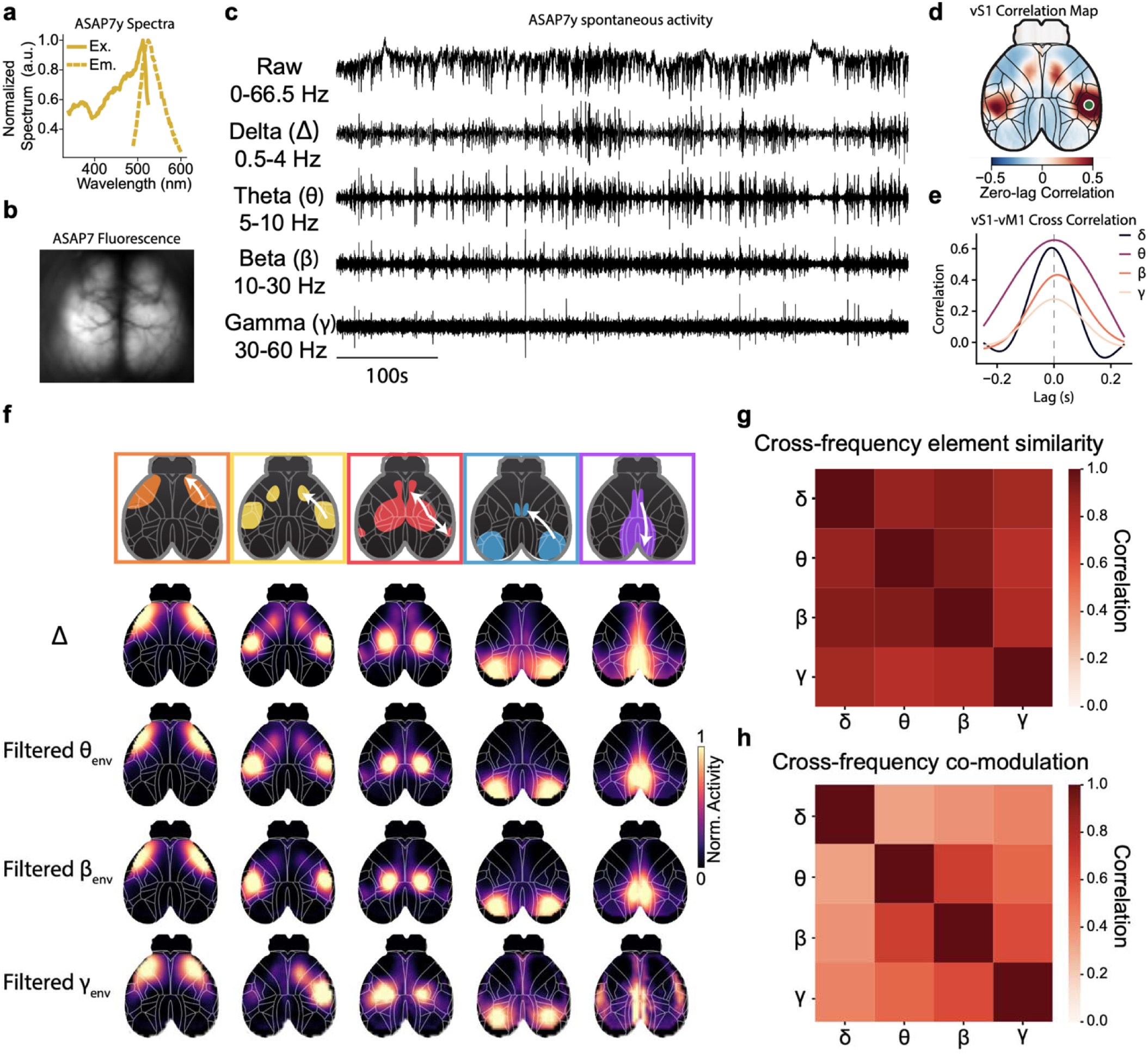
Voltage imaging reveals shared directional element structure across frequency bands. **a.** Excitation and emission spectra of ASAP7y. **b.** Widefield ASAP7y fluorescence imaged through a cleared skull preparation. **c.** Example ASAP7 voltage traces bandpass-filtered into canonical frequency bands. **d.** Correlation map between broadband vS1 voltage activity and the rest of the cortex, revealing the active sensory (AS) network. **e.** Cross-correlation between AS nodes showing consistent but modest lead of vS1 activity across frequency bands. Single-replicate example data. **f.** Frequency-resolved directional elements. Applying the directional element discovery pipeline to band-limited voltage signals reveals spatial motifs that recapitulate those identified in Ca² imaging across all frequency bands. **g.** Cross-frequency element similarity. Pairwise correlations between matched directional elements across frequency bands demonstrate conservation of spatiotemporal structure. **h.** Cross-frequency temporal co-modulation. Temporal loadings of frequency-specific elements show correlated fluctuations across bands, with delta (δ)-band activity exhibiting progressively stronger coupling toward higher-frequency gamma (γ)-band dynamics.

Expression of ASAP7y across dorsal cortex by whole-brain AAV delivery produced high-quality fluorescence images comparable to those obtained with modern GCaMP indicators **(Fig. 5b)**. Single-session traces from vS1 revealed rich broadband activity that could be decomposed into distinct frequency bands—delta, theta, beta, and gamma—each potentially capturing different aspects of the underlying neural dynamics **(Fig. 5c)**^52,53^. Seed-based correlation analysis of vS1 delta-band activity revealed a directional correlational structure closely mirroring that observed with Ca² imaging, with strong homotopic contralateral coupling and recruitment of vM1 **(Fig. 5d)**. Temporal cross-correlation between vS1 and vM1 again demonstrated a consistent lag structure **(Fig. 5e)** across all frequency bands except beta, confirming that the directional propagation relationships identified with Ca² imaging were preserved in voltage signals.

Applying the network element discovery pipeline to the delta-band filtered voltage imaging data yielded a set of spatiotemporal directional elements with striking correspondence to those identified with Ca² imaging **(Fig. 5f, Extended Data Fig. 9, Supplementary Video 5)**, recapitulating the AL, AS, MS, V, and MA organizational structure in this independent modality. Extending the analysis to theta, beta, and gamma bands revealed that directional elements with recognizable spatial organization could be recovered across all frequency bands examined, though with band-specific differences in spatial extent and the sharpness of network boundaries **(Fig. 5f)**^52^. For example, higher frequency bands produced stronger source-region loadings in AS, MS, and (to a lesser extent) V elements, whereas in the gamma band, AL showed a bias towards sink activity.

To quantify cross-band correspondence, we computed pairwise cross-correlations among matched directional elements across frequency bands, mirroring the cross-cell-type analysis of **Fig. 1j**. This analysis revealed high similarity among matched elements across delta, theta, beta, and gamma bands for the majority of directional element pairs **(Fig. 5g)**, indicating that the large-scale spatial organization of cortical directional elements is a broadband phenomenon not limited to the slower fluctuations captured by Ca² imaging. To further characterize the relationship between frequency bands, we compared temporal loadings of matched directional elements across bands. We found that temporal loadings were broadly correlated across frequencies, with a systematic gradient in which delta-band loadings showed progressively stronger correspondence with higher frequency bands **(Fig. 5h),** consistent with the observation that slow fluctuations in band-limited power are coherently expressed across frequency bands in primate cortex^54^. Together, these results demonstrate that the directional element framework generalizes across both indicator modality and frequency band spanning the full range of relevant neural timescales.

## Discussion

We developed an approach to record spontaneous cortex-wide activity, without scanning or tiling, over many tens of hours, and applied this method to unbiased screening for regularities allowing for temporal delays in signal propagation. This resulted in extraction of a set of directional elements: consistent spatiotemporal patterns of cortical activity that generalized across cell types and timescales with demonstrable relevance to behavior. The results establish a comprehensive framework for understanding the structure, behavioral embedding, and causal control principles of large-scale cortical network organization in the mammalian brain.

The conservation of directional elements across excitatory and inhibitory cell types was among the most striking initial findings. Given the well-documented distinct physiological properties and computational roles of VIP, Sst, and pyramidal neuron populations, one might have expected each cell class to exhibit a unique large-scale organizational signature. Instead, any of the four cell-type-specific basis sets comparably explained spontaneous activity variance in other cell types as well. This result may reflect a fundamental property of cortical mesoscale organization: at the spatial and temporal scales sampled by widefield imaging, network membership is determined by neuron positioning rather than by cell type. This finding of course does not preclude cell-type-specific operations at finer scales— indeed, the well-established roles of VIP-mediated disinhibition and Sst-mediated gain control almost certainly shape the within-network computations that these elements perform— but suggests that these operations are coordinated across cell types in a way that preserves the collective spatial structure of network activation.

The source-to-sink control principle established by the all-optical control assay provides a mechanistic account of how these directional elements are recruited. Across all five neocortical directional elements, brief optogenetic activation of early-active source regions sufficed to drive downstream network-wide activity, while stimulation of late-active sink regions produced only local responses. This asymmetry may represent a general form of findings from individual sensory systems—in which silencing primary sensory areas abolishes downstream responses while manipulating higher-order areas has modest retrograde effects^55–57^— now extended to motor^58^ and associative systems including the MA element. Importantly, by grounding the definition of source and sink regions in the intrinsic spatiotemporal dynamics of spontaneous activity rather than sensory-evoked responses, this new approach is agnostic to the specific sensory modality or behavioral context, enabling principled investigation of networks like MA that cannot be directly assayed by peripheral stimulation. These findings raise the question of what connectivity motifs give rise to the source-to-sink asymmetry — a question that recurrent network models of sequential activity generation may be well positioned to address^59,60^.

Pharmacological challenges coupled with imaging and perturbation assays revealed a striking dissociation between two aspects of network organization. Spontaneous directional elements event rates were profoundly and distinctively reorganized by different major drug classes—so reliably that drug condition could be decoded from element dynamics with high accuracy within minutes of injection. Yet despite these large perturbations of spontaneous dynamics, the ability of source regions to causally recruit downstream activity was preserved across all conditions. This dissociation suggests that the source-to-sink control hierarchy reflects stable structural configuration rather than a contingent feature of any particular dynamical state, and raises the intriguing possibility that source-targeted stimulation could restore normal patterns of network activity even when spontaneous dynamics are severely disrupted—a potentially important consideration for therapeutic neuromodulation.

Visual detection behavioral experiments combined with all-optical recording and control of network state demonstrate that these network dynamics are not merely correlates of behavior but causally shape behavior. Pre-stimulus optogenetic recruitment of AL, AS, and MA sources each improved visual detection accuracy and reduced reaction time, while Visual cortex stimulation had no effect, and the specific role of MA network engagement in quantitatively tuning detection performance was particularly noteworthy. The MA element exhibits certain functional and anatomical similarities to the human default mode network (which has been conventionally associated with internally directed cognition and has been hypothesized to compete with external sensory processing) but our results argue against a simple correspondence; notably, pre-stimulus MA recruitment was consistently facilitatory, and moreover on a trial-by-trial basis we found that stronger optogenetically-evoked MA activity predicted better detection outcomes. We suggest that in the context of a learned detection behavior, MA network engagement reflects the activation of an internalized action-outcome model that prepares the animal to respond appropriately to the expected stimulus—a form of predictive, model-based processing that actively supports rather than competes with perception^61,62^. These findings may provide some of the clearest causal evidence to date for a reframing of default mode network-related components away from passive mind-wandering and toward an active role in situational model construction and behavioral preparation.

Taken together, these findings establish several principles of large-scale cortical organization with broad relevance. First, the directional elements provide a cell-type-agnostic, spatiotemporally resolved reference framework for cortical dynamics that may serve as a common language for comparing results across experimental paradigms, brain states, and pharmacological conditions. Second, the source-to-sink hierarchy provides a principled map of where in a network causal interventions may be effective^63^—a consideration directly relevant to the design of closed-loop neuromodulation strategies. Third, drug decoding establishes directional element event rates as a sensitive translational biomarker of pharmacologically modulated brain state, potentially bridging preclinical and clinical assessments of drug target engagement. Fourth, large-scale directional element state causally shapes perceptual behavior; in particular, stronger trial-by-trial causal MA network engagement consistently predicted successful response behaviors—revealing that the medial associative network, conventionally associated with internal cognition, actively facilitates rather than competes with sensory detection. Fifth, preservation of directional elements across voltage imaging and across spectral frequency bands—together with broad temporal co-expression across frequencies—establishes robust, modality-invariant measurement of cortical organization that spans the full spectrum of neural activity. Together, the directional elements emerging from unbiased screening establish a detailed and direct link among between local dynamics, brainwide activity, and behavior.

## Acknowledgements

We thank I. Kauvar, T.A. Machado, S. Vesuna, S. Quirin, T.X. Liu, and all members of the Deisseroth lab, as well as S. Ganguli, I. Landau, M. Bilokur, H.S. Hunt, and all members of the Ganguli lab for input on the project and feedback on the manuscript. This work was supported by the National Institutes of Health grant U19NS118284 (K.D.), National Institutes of Health grant P50DA042012 (to K.D.), National Institutes of Health grant R01 MH086373 (K.D.), the Gatsby Foundation (K.D.), a Knight Initiative for Brain Resilience Catalyst Award (K.D.), the Keck Foundation (K.D.), R01 EY022638 (to T.R.C.), P30EY026877 (to T.R.C.), The Chan Zuckerburg Biohub, San Francisco (to T.R.C.), a Stanford Bio-X Seed Grant (to T.R.C.). UM1MH136462 (to M.Z.L.) and 1RM1NS132981 (to M.Z.L.). Y.W. is a Howard Hughes Medical Institute Fellow of The Jane Coffin Childs Memorial Fund. Y.A.H. was supported by the Stanford BioX Bowes Fellowship.

## Author Contributions

J.K., A.D.W., Y.W., A.M., and K.D. designed the study. J.K. built the widefield macroscope, all-optical stimulation system, and visual detection task apparatus. J.K., K.S., C.K., A.M., Y.W., A.D.W., W.W., and M.H. collected the data. J.K. analyzed the experimental data, with assistance from A.D.W., A.M., and S.D. in analysis conceptualization. Y.A.H., T.C., and M.L. developed ASAP7y. C.R., K.E.P., and N.A. cloned all viral vectors. J.K., A.D.W., Y.W., and K.D. wrote the manuscript with input from all authors. K.D. supervised all aspects of the project.

## Code and Data Availability

Processed data files including neural recordings, behavior movies, and Bpod behavioral session files used for all analyses will be made available as a repository on Zenodo. The raw widefield imaging datasets generated during this study will also be available from the corresponding author upon request. Custom Python scripts for data acquisition, preprocessing, and analysis will be deposited in a public repository upon publication.

---

**Extended Data Figure 1.**
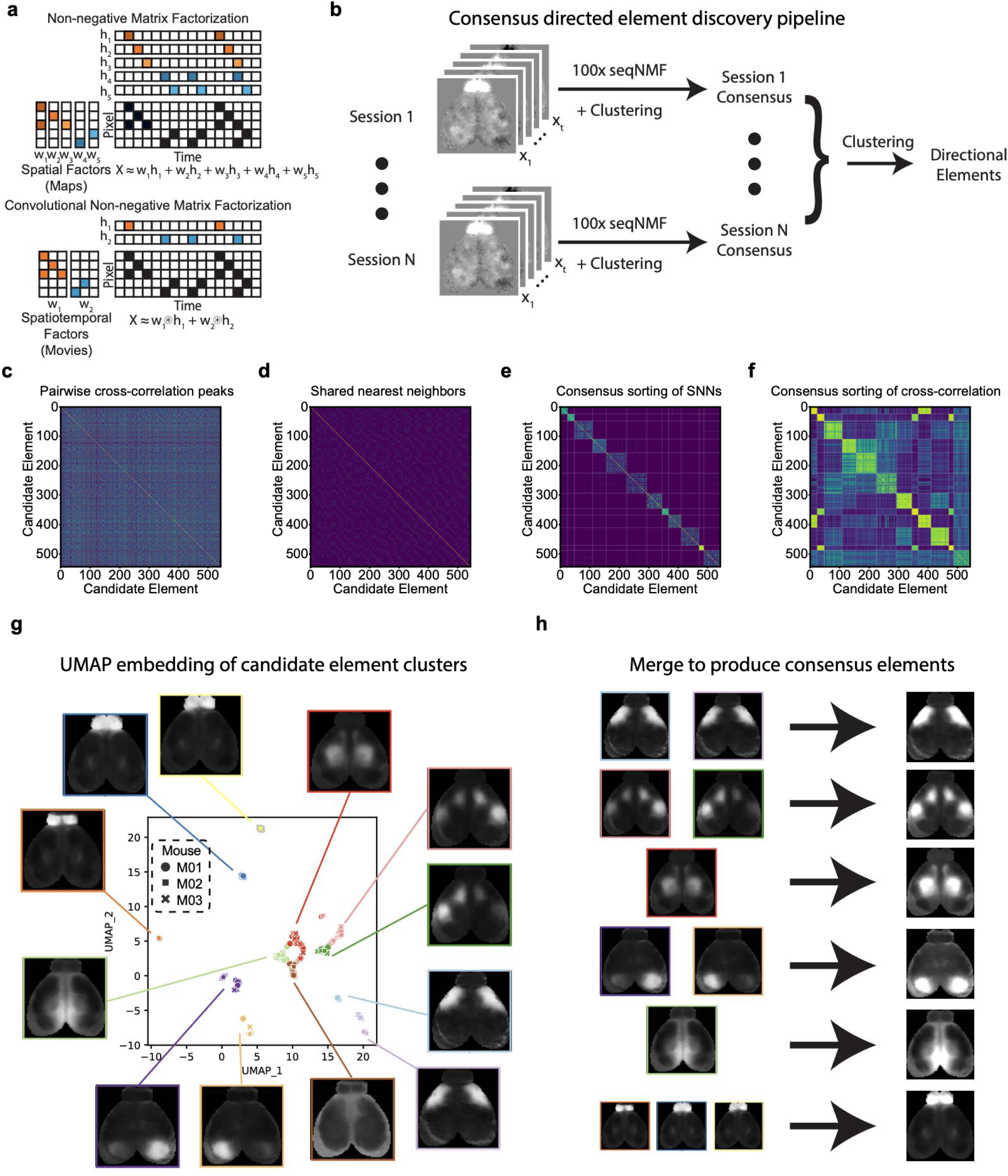
Directional element discovery pipeline. **a.** Comparison of matrix factorization approaches. Comparison of standard (top) and convolutional (bottom) NMF decomposition of synthetic widefield Ca^2+^ data. Both methods factorize the pixels-by-time data matrix into a lower-dimensional set of spatial (or spatiotemporal) and temporal components; however, standard NMF requires more factors to capture the temporal dependencies in the data. **b.** Consensus clustering for directional element discovery. Individual sessions are factorized one hundred times with seqNMF, then the resulting candidate elements are clustered based on their pairwise cross-correlation peaks, temporally aligned, and averaged to produce session-level consensus elements. The session-level consensus elements from different recordings are then clustered in the same fashion, resulting in the final set of directional elements. **c**–**f)** Intermediate outputs of the clustering pipeline applied to all Cux2 candidate elements. **c)** Pairwise cross-correlation peak matrix between candidate directional elements. **d)** Shared Nearest Neighbor (SNN) graph derived from pairwise correlations. **e)** SNN matrix after consensus clustering and sorting reveals block structure corresponding to candidate element clusters. White lines outline borders of identified clusters. **f)** Cross-correlation matrix after consensus sorting, confirming the same cluster structure and highlighting strong cross-block correlations among apparently similar clusters. **g)** Uniform Manifold Approximation and Projection (UMAP) embedding of all Cux2 candidate elements, color-coded by cluster identity and with symbol denoting mouse of origin. Spatial maps display the consensus directional elements derived from each cluster. **h)** Final merging of consensus elements: duplicate candidate elements arising from the same network are aligned in time and averaged to produce the final consensus directional elements.

**Extended Data Figure 2.**
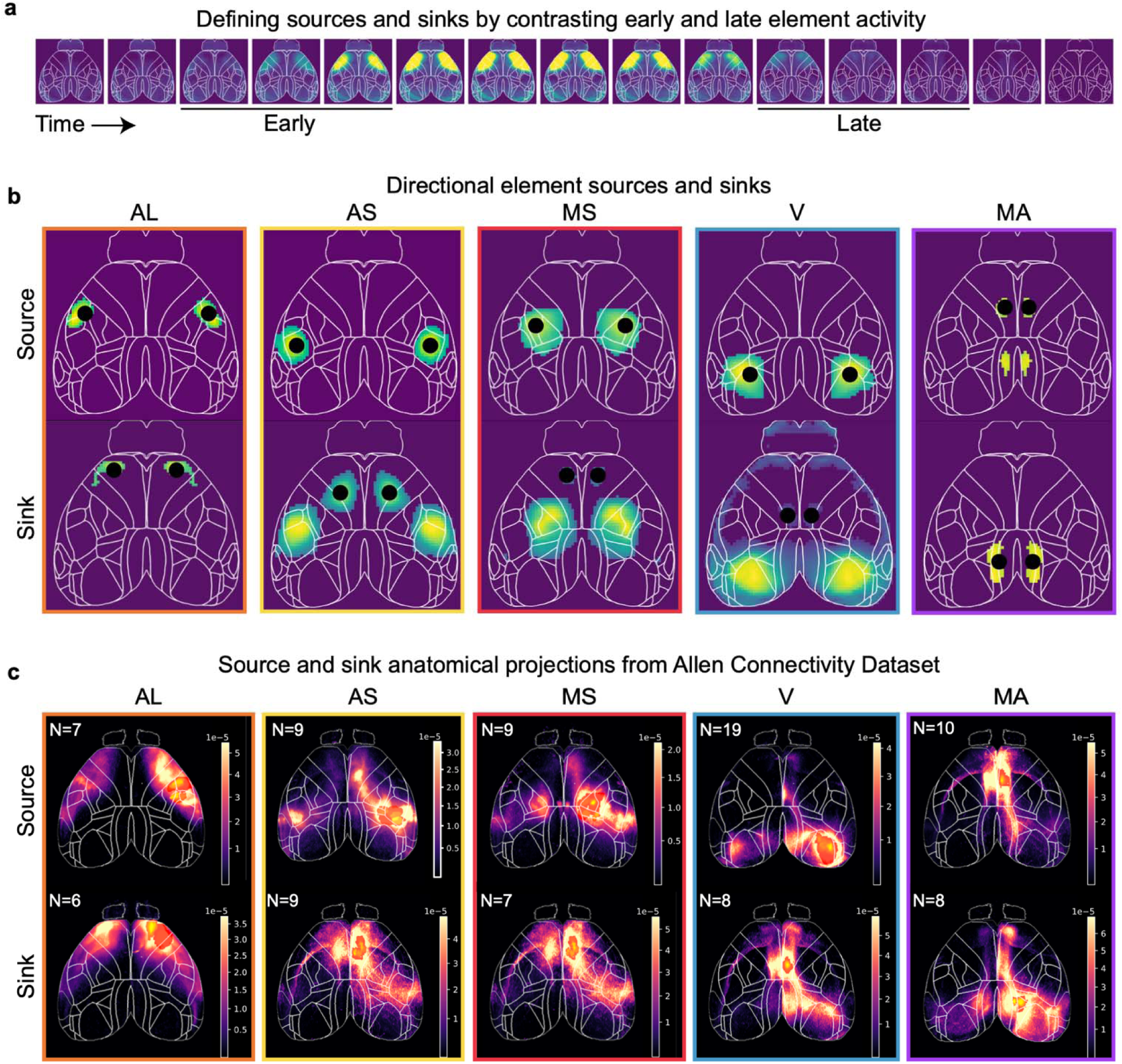
Source/sink identification and analysis of anatomical connectivity. **a.** Defining source and sink regions from directional element spatiotemporal dynamics. Frames spanning the early and late phases of each element’s activity movie are contrasted to identify regions of early (source) and late (sink) activation. **b.** Source (top) and sink (bottom) spatial maps for each of the five major neocortical directional elements: Anterolateral (AL, orange), Active Sensory (AS, yellow), Medial Somatomotor (MS, red), Visual (V, blue), and Medial Associative (MA, purple). Maps are overlaid on the Allen Brain Atlas cortical outline^64^. Black circles denote the center of mass of source or sink activity.**c.** Analysis of anatomical connections from Allen Connectivity Dataset. Average source (top) and sink (bottom) projection maps across all directional elements. N values indicate the number of individual experiments contributing to each average. Maps reveal strong concordance between long range projection patterns and identified functional directional elements. Notably, projections show no evidence of directional bias between source and sink.

**Extended Data Figure 3.**
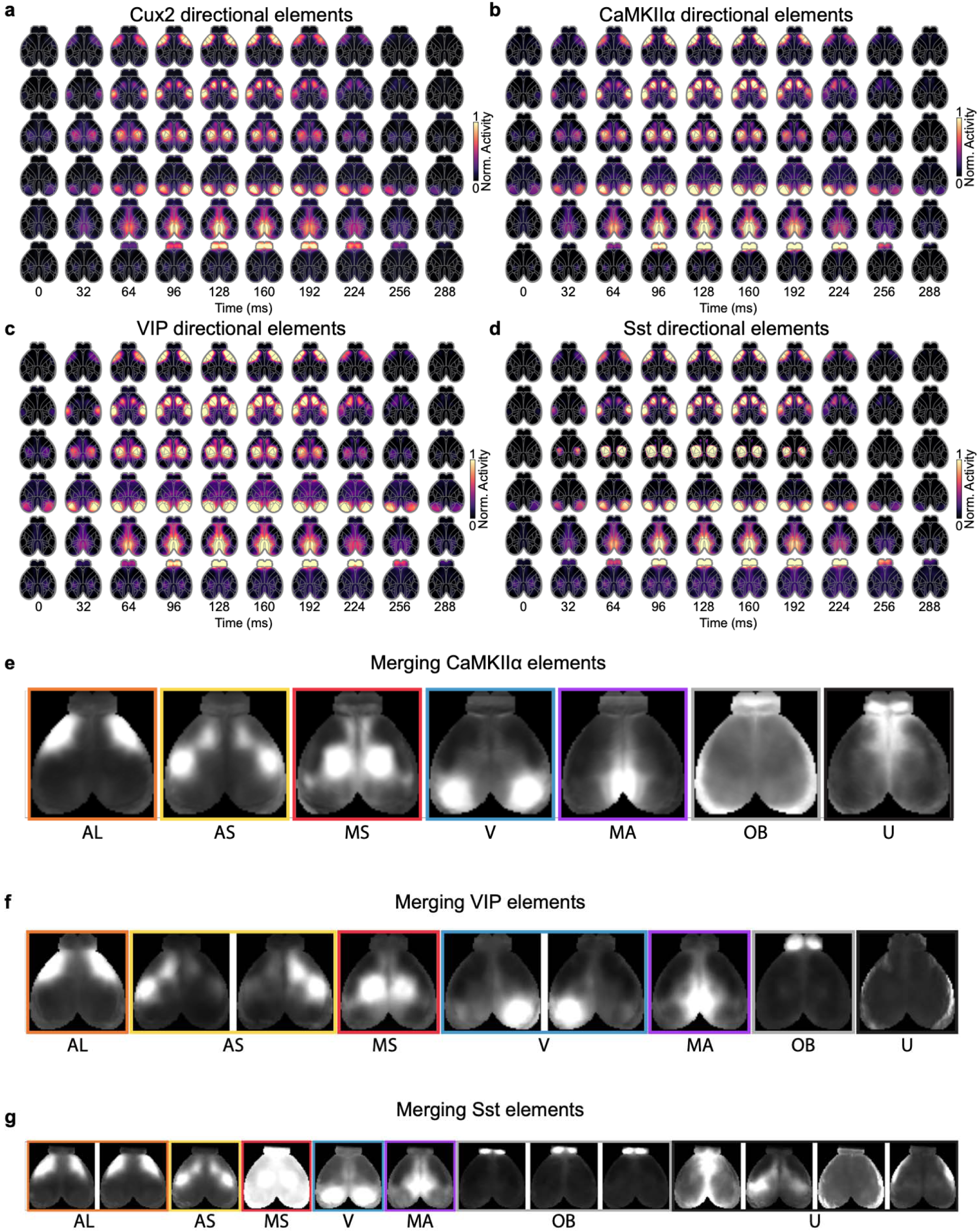
**Directional elements identified across cell type-specific imaging cohorts**. **a–d,** All cell-type specific directional elements: a, Cux2 (layer 2/3 excitatory neurons), b, CaMKIIα (broad excitatory population), c, Vasopressin Intestinal Peptide (VIP) interneurons, and d, Somatostatin (Sst) interneurons. Each panel shows spatiotemporal movies of all recovered elements displayed as a sequence of frames (0–288 ms; color scale: normalized activity 0–1). The same set of directional element types is recovered across all cell-type datasets. **e–g,** Raw consensus directional elements from each cell-type-specific cohort, grouped by hand-selected merges to produce final directional element sets: e, CaMKIIα, f, VIP, and g, Sst. AL: Anterolateral, AS: Active Sensory, MS: Medial Somatomotor, V: Visual, MA: Medial Associative, OB: Olfactory, U: Uncategorized.

**Extended Data Figure 4.**
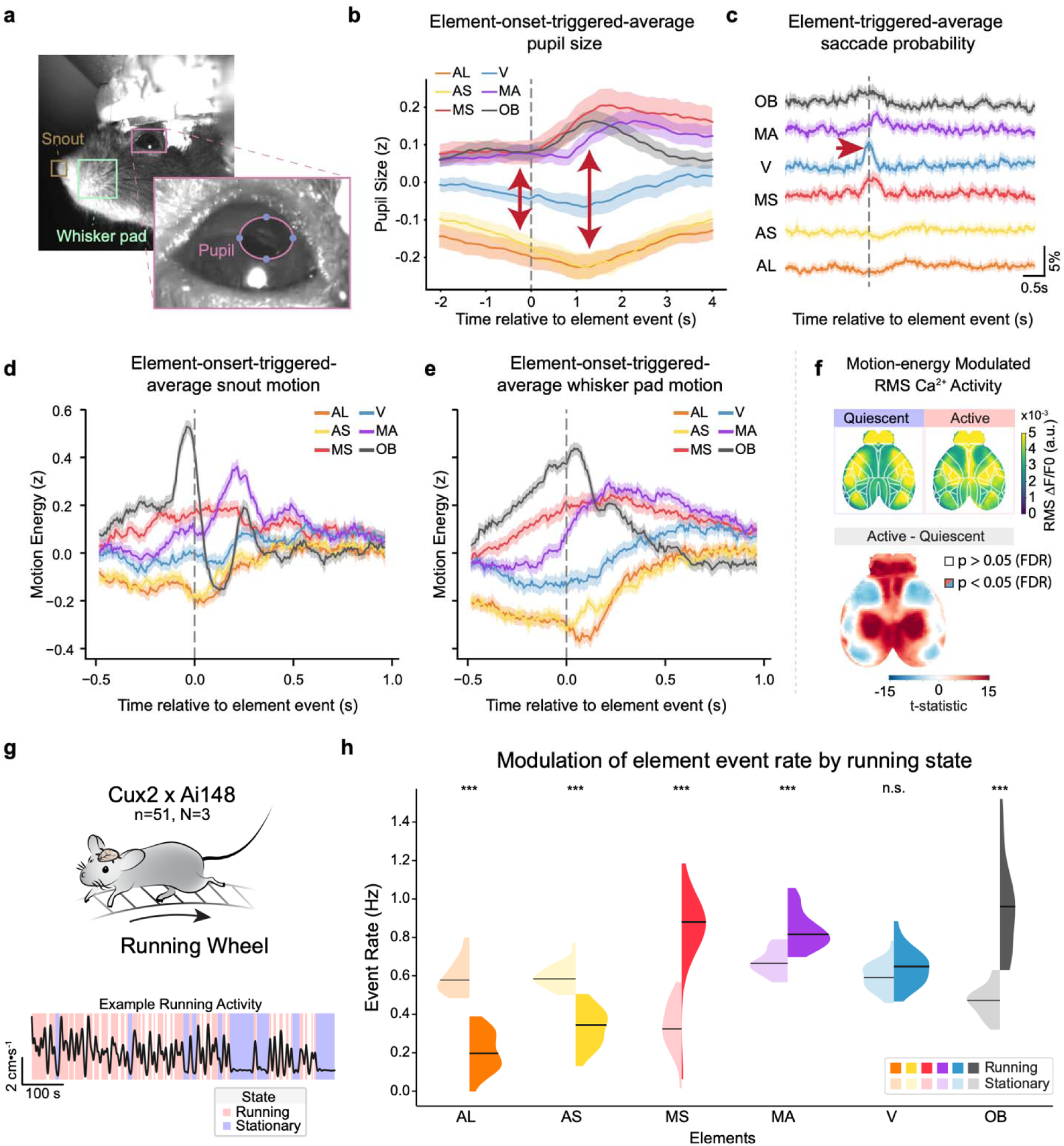
Behavioral correlates of directional element events. **a.** Facial video monitoring during widefield imaging. Regions of interest over the snout (brown) and whisker pad (green) were extracted and total motion energy computed. The pupil was tracked by using DeepLabCut to identify four keypoints—top, bottom, nasal, temporal—to which an ellipse was fit to determine pupil size and position. **b.** Element-triggered average pupil size aligned to directional element event onset (dashed line at time=0) for each element. AL and AS directional element events were observed when pupil was relatively small, and were followed by further constriction, whereas MS, MA, and OB showed the opposite pattern, with V showing intermediate dynamics between these two extremes. Shading: 95% CI. **c.** Element-triggered average saccade probability for each directional element. V shows a clear bump in saccade probability centered at event onset (time = 0). Scale bar: 5% probability, 0.5s. Shading: 95% CI. **d.** Element-triggered average normalized snout motion energy aligned to directional element event onsets. OB element showed a striking ∼3.35 Hz oscillatory waveform in snout motion energy starting approximately 120 ms prior to, and peaking ∼30ms before, onset of the cortical event, consistent with the known coupling between rhythmic sniffing behavior and olfactory bulb activity. **e.** Element-triggered average normalized whisker pad motion energy aligned to element event onsets. OB was again distinctive, showing a sustained ramp in whisker pad motion beginning ∼500 ms before event onset and peaking sharply at event time, suggesting that coordinated orofacial exploratory behavior—involving both rhythmic sniffing and whisker protraction—precedes and likely drives olfactory cortical recruitment. MS and MA events occurred at times of elevated baseline whisker pad motion, while AL and AS events were associated with relatively low whisker pad motion energy at event onset, consistent with their preferential occurrence during quiescent behavioral states. **f.** Statistical analysis of state-dependent RMS Ca^2+^activity differences. A pixelwise paired t-test across Active and Quiescent states was performed and p-values FDR corrected. A map of the t-statistic, masked to only include significant p-values (p<0.05 after correction) depicts areas of significant relative increases in RMS in the Active (warm colors) and Quiescent (cool colors) states. **g.** Experimental setup for running modulation analysis. Head-fixed Cux2×Ai148 mice (N=3 mice, n=51 sessions, 10.2 h) were imaged on a running wheel while locomotion was monitored. Example running trace shown (black), with running (red) and stationary (blue) epochs highlighted. **h.** Directional event rate modulation by running behavior. Mirroring the analysis of orofacial motion energy (Fig 1), AL and AS event rates were suppressed during running bouts, whereas MS, MA, and OB rates were enhanced (Paired *t*-test per element, *** = *p* < 0.001, n.s., not significant, N=3, n=51).

**Extended Data Figure 5.**
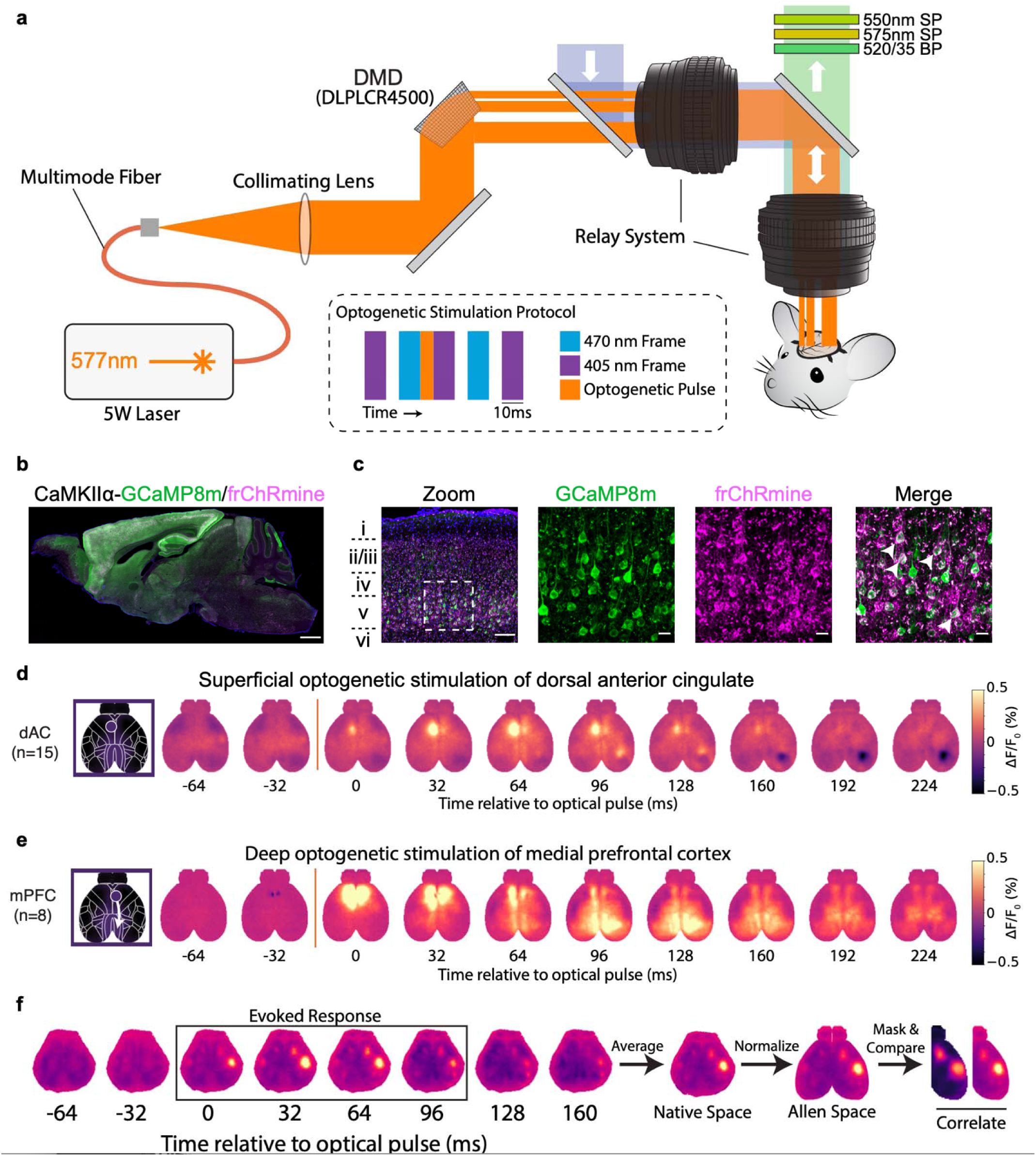
Patterned optogenetic stimulation and simultaneous widefield Ca^2+^ imaging. **a.** Schematic of the combined optogenetic stimulation and widefield imaging system. A 577 nm laser is coupled via multi-mode fiber into a digital micromirror device (DMD; DLPLCR4500) for patterned illumination. Light is directed through a lens relay system onto the dorsal cortex of a head-fixed mouse. Emission is collected through shortpass (550 nm SP, 575 nm SP) and bandpass (520/35 BP) filters. The optogenetic stimulation protocol (inset) interleaves 405 nm (isosbestic) and 470 nm (GCaMP excitation) imaging frames (10 ms) with 577 nm optogenetic pulses (6 ms) delivered between frame exposures. **b.** Sagittal section of a mouse co-expressing GCaMP8m (green) and frChRmine (magenta) under the CaMKIIα promoter, confirming dual viral expression across cortical layers. Scale bar: 1 mm. **c.** High-magnification confocal images showing individual channels (GCaMP8m, frChRmine) and the result of merging. White arrowheads mark cells co-expressing both constructs. Approximate cortical layer designations (i–vi) shown at left. Scale bar: 20 µm. **d.** Mean widefield fluorescence response (ΔF/F, %) to superficial optogenetic stimulation targeting the dorsal anterior cingulate cortex (dAC; n=15 replicates), shown as sequential frames from −64 to +224 ms relative to pulse onset (dashed line). Activation spread from the stimulation site locally and contralaterally, but did not produce a noticeable wave. **e.** As in d, for deep optogenetic stimulation targeting the medial prefrontal cortex (mPFC; n=8 replicates). A striking wave propagating posteriorly along the midline into RSC and outward into parietal cortices was observed. **f.** Workflow for computing the stimulation-evoked response used for comparison with directional element spatial maps. Frames corresponding to the evoked response window were averaged, transformed to Allen CCFv3 space, masked to contain only the stimulated hemisphere, then spatially correlated with the maximum intensity projection of the corresponding directional element.

**Extended Data Figure 6.**
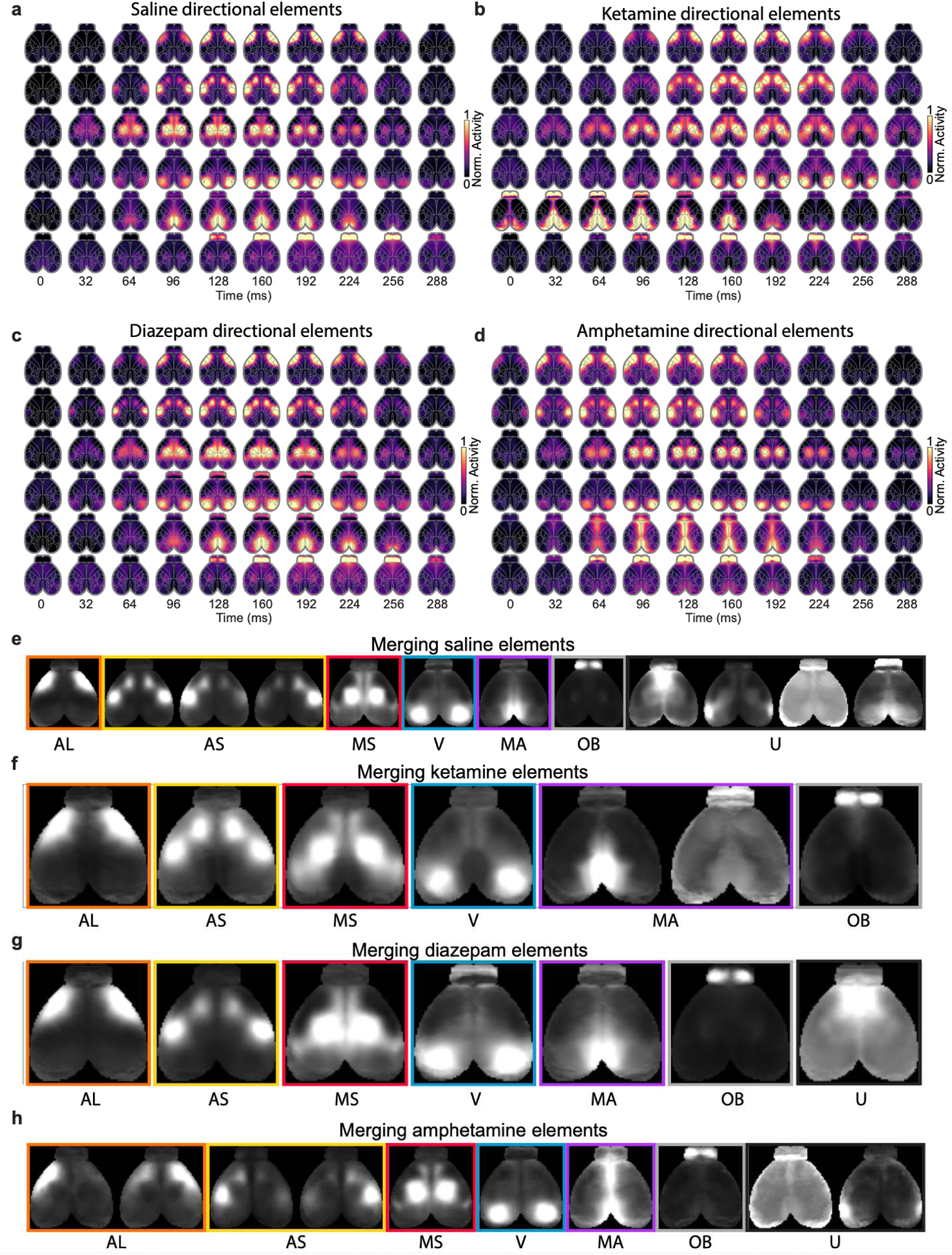
Drug-specific directional elements. **a–d)** All directional elements identified across pharmacological conditions: a, saline control, b, ketamine, c, diazepam, and d, amphetamine. Each panel displays the full set of spatiotemporal directional element movies as sequential frames (0–288 ms; color scale: normalized activity 0–1). The same six major elements (AL, AS, MS, V, MA, OB) are broadly recovered across conditions. **e–h)** Raw consensus directional elements for each compound showing chosen merges, for e, saline, f, ketamine, g, diazepam, and h, amphetamine. Elements are labeled by type.

**Extended Data Figure 7.**
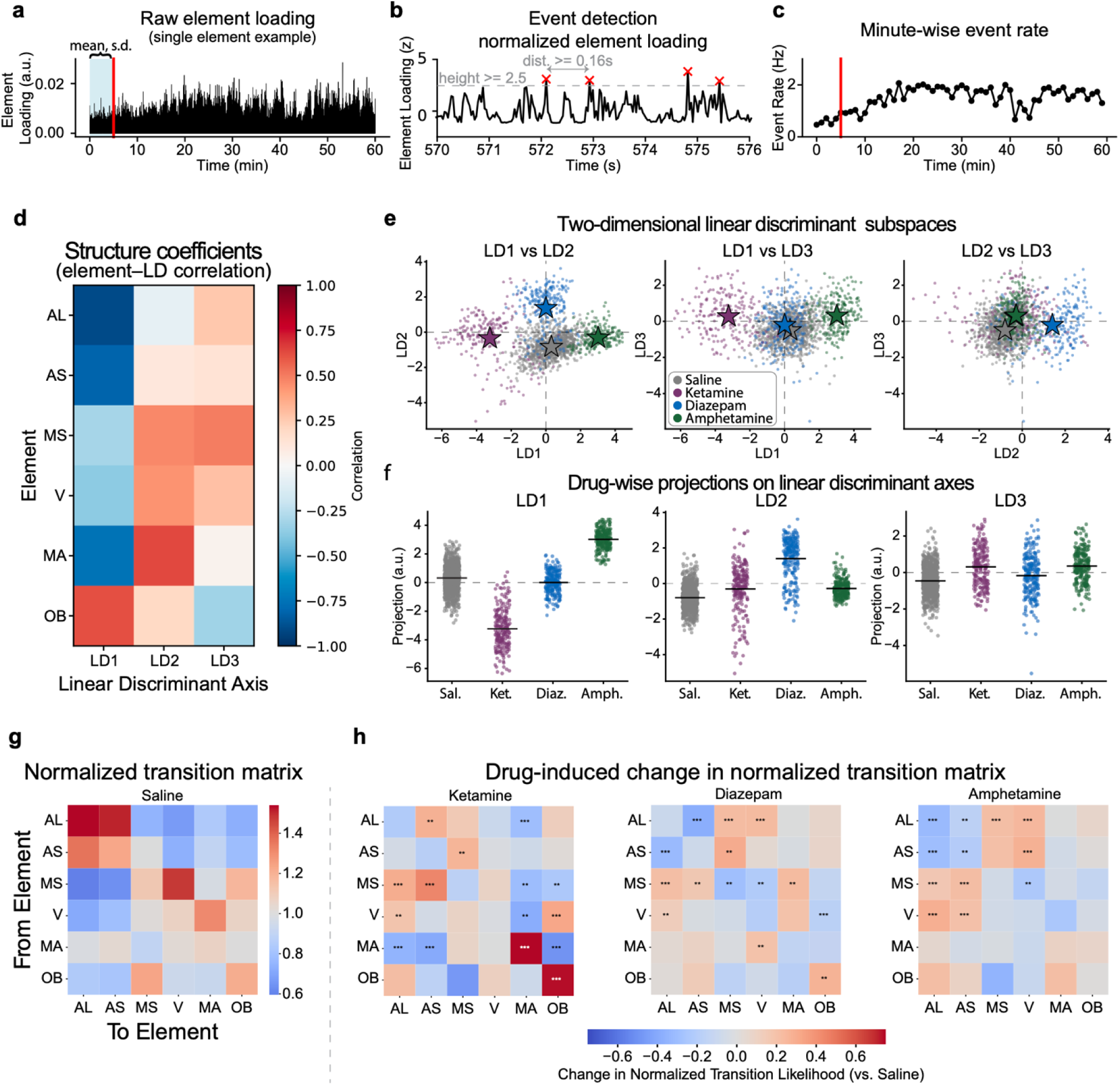
Pharmacological effects on directional element dynamics and state transitions. **a.** Example raw element loading timeseries for a single directional element over a 60-minute recording session. Red line at 5 minutes indicates injection time; mean and standard deviation from the initial 5-minute baseline period used for normalization. **b.** Zoomed view of the normalized directional element loading trace showing the event detection procedure. Peaks exceeding a z-score threshold of 2.5 and separated by at least 0.16 s are marked as events (red x’s). **c.** Minute-wise element event rate over the session. Data are divided into 1-minute bins and events counted and converted to an event rate (Hz). **d.** Heatmap of structure coefficients (element–linear discriminant correlations) relating each directional element type (AL, AS, MS, V, MA, OB) to the three linear discriminant axes (LD1–LD3) derived from pharmacological condition classification. Warm colors indicate positive correlation; cool colors indicate negative correlation. **e.** Two-dimensional projections of session-level element activity onto pairs of linear discriminant axes (LD1 vs. LD2, LD1 vs. LD3, LD2 vs. LD3). Each point represents one minute of data, color-coded by drug condition (saline: gray; ketamine: cyan; diazepam: magenta; amphetamine: olive). Stars mark condition centroids. Drug conditions are well-separated in LD space. **f.** Distribution of projections onto each linear discriminant axis (LD1–LD3), separated by drug condition. Horizontal lines indicate condition means. Ketamine and amphetamine are strongly separated along LD1; diazepam is separated from other compounds along LD2. **g.** Normalized element-to-element transition matrix under saline conditions. Values indicate the relative likelihood of transitioning from one directional element type (rows) to another (columns), normalized by each element’s marginal event rate. Diagonal entries reflect self-transitions. **h.** Drug-induced changes in the normalized transition matrix relative to saline, shown separately for ketamine, diazepam, and amphetamine. Color scale: change in normalized transition likelihood (blue: decreased, red: increased). Asterisks indicate statistically significant changes (permutation test performed by shuffling event times among directional elements 1000 times, ** p<0.01, *** p<0.001).

**Extended Data Figure 8.**
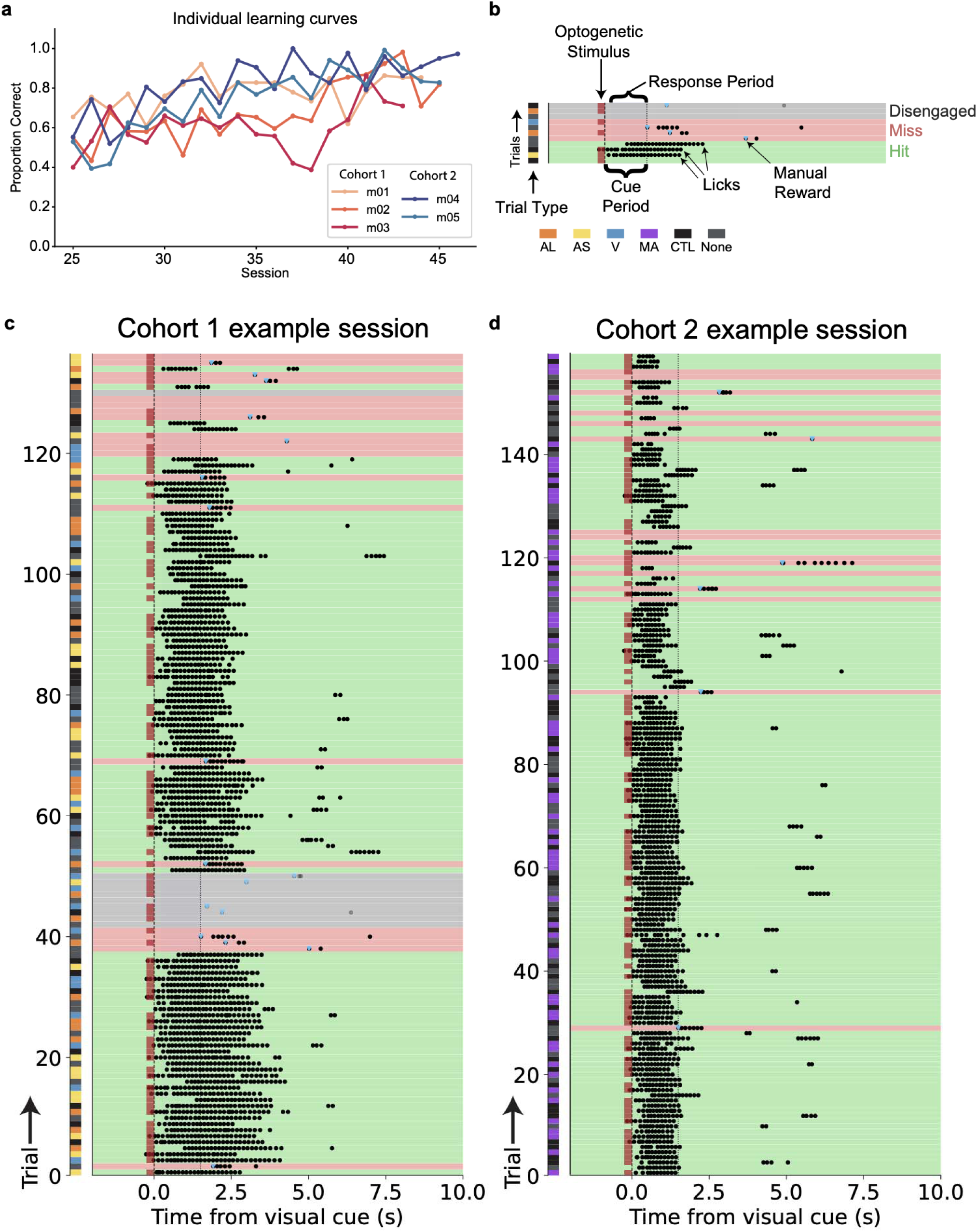
Behavioral task design and performance for visual detection task. **a.** Individual learning curves for all mice across sessions, showing the proportion of correct trials as a function of session number. Two training cohorts are shown: Cohort 1 (m01–m03, warm colors) and Cohort 2 (m04–m05, cool colors). **b.** Schematic of the lick raster highlighting task structure and trial classification. Each row is a trial. Mice reported cue detection by licking during the response period; hits (correct licks, green), misses (no lick, pink), and disengaged trials (5 ore more consecutive misses, gray) were distinguished. Experimenter occasionally delivered manual rewards (blue triangle) to encourage engagement. On a random subset (∼2/3) of trials, a brief optogenetic stimulus was delivered immediately prior to stimulus onset. Trial types color-coded by the targeted directional element (AL, AS, V, MA, Control, and No Stimulation**). c.** Lick raster for an example Cohort 1 session. Each row is a trial, sorted by trial number. The color bar on the left indicates trial type. Black dots show individual lick times; blue dots indicate rewarded licks; the red dashed line marks stimulus onset and the dotted line indicates the end of the response window. Shading denotes trial outcome (green: hit, pink: miss, gray: disengaged). **d.** As in **c**, for an example Cohort 2 session (MA element stimulation only).

**Extended Data Figure 9.**
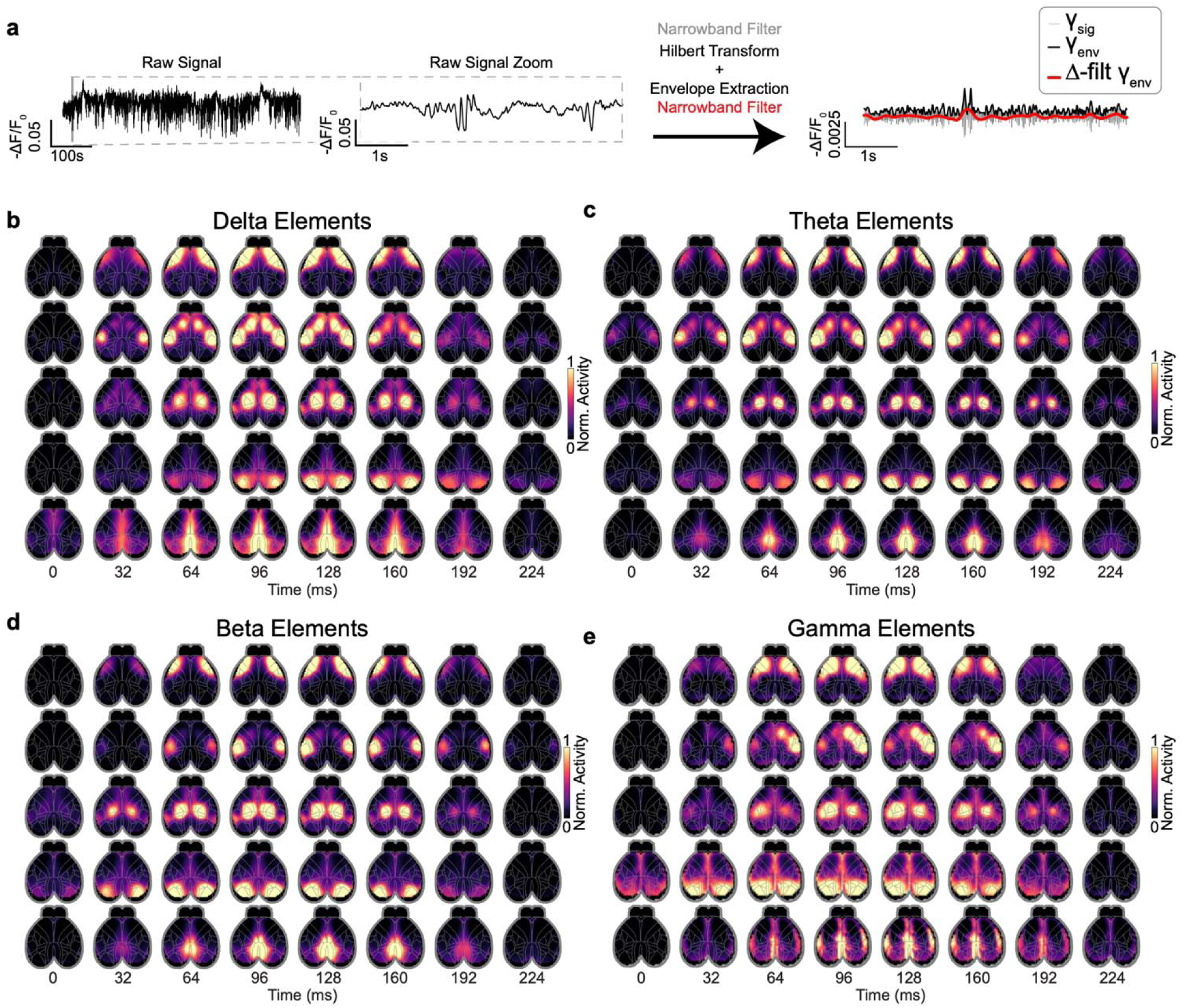
Frequency band-specific directional element discovery. **a.** Signal processing pipeline for extracting frequency-band-specific ASAP7 signals. The raw widefield ΔF/F trace (left) was bandpass filtered into the frequency band of interest (narrowband filter), and the analytic envelope was extracted using the Hilbert transform (right). The resulting signal (γ_env, black) and its narrowband-filtered version (Δ-filt γ_env, red) were shown alongside the original broadband signal (γ_sig, gray).**b–e,** All directional elements recovered by applying the discovery pipeline to envelope signals in four canonical frequency bands: b, delta, c, theta, d, beta, and e, gamma. Each panel displays the full set of spatiotemporal directional element movies as sequential frames (color scale: normalized activity 0–1). The same core directional element types were recovered across all frequency bands, demonstrating that the directional element structure was not specific to Ca^2+^ activity nor a specific spectral regime.

**Supplementary Video 1: Cortex-wide spontaneous activity displays rich spatiotemporal dynamics.**

Example recording of Cux2::GCaMP6f across the entire dorsal cortex, normalized to Allen CCFv3, with simultaneous orofacial video. Video is played at speed of brain imaging (31.25 fps). Orofacial video recorded at 125fps and temporally downsampled to match brain imaging. Related to data in Figure 1.

**Supplementary Video 2: A common set of directional elements is shared across cell types and layers.**

Cell type-specific directional elements derived from Cux2, CaMKIIα, VIP, and SST datasets show remarkable similarity. Playback slowed to 1/4x speed (7.8 fps). Related to data in Figure 1.

**Supplementary Video 3: Source stimulation recruits directional elements.**

Optogenetic stimulation of source, but not sink, sites in mice expressing GCaMP8m and frChRmine under the control of the CaMKIIα promoter is sufficient to recruit downstream directional element sites. Blue wavy arrows denote the stimulated site; green arrows indicate downstream recruitment. Playback slowed to 1/4x speed (7.8 fps). Related to data in Figure 2.

**Supplementary Video 4: Directional elements are conserved across pharmacological manipulations.**

Directional elements derived from the peak drug phase (5-35 min) following administration of saline, ketamine, diazepam, or amphetamine show conserved structure. Playback slowed to 1/4x speed (7.8 fps). Related to data in Figure 3.

**Supplementary Video 5: Directional elements are recapitulated by voltage imaging and conserved across frequency bands.**

Voltage imaging of spontaneous cortex-wide ASAP7y activity followed by frequency-specific directional element discovery recovers the same set of elements observed with Ca2+ imaging. Playback slowed to 1/4x speed (7.8 fps). Related to data in Figure 5.

## Methods

### Animals

All procedures were conducted in accordance with protocols approved by the Stanford University Institutional Animal Care and Use Committee (IACUC) and National Institutes of Health guidelines. Both male and female C57BL/6J-background mice aged 8–10 weeks at surgery were used. Transgenic lines used were: Cux2-CreER^T2^ (MGI 5014172), Sst-Cre (JAX 013044), VIP-Cre (JAX 010908), Ai148 (JAX 030328), and C57BL/6J (JAX 000664). Mice homozygous for Ai148 and heterozygous for the Cre transgenes from each driver line were bred to produce double transgenic mice with the genotypes Cux2-CreER;Ai148, VIP-Cre;Ai148, and SSt-Cre;Ai148.

### Transduction and Viral Vectors

GCaMP expression in Cux2-CreER;Ai148 mice was induced with three intraperitoneal injections of 100 mg·kg^-1^ tamoxifen over five days. For all-optical experiments, 4-week-old C57BL/6J mice received a 100 µL retro-orbital injection containing a mixture of AAV-PHP.eB-CaMKIIα-GCaMP8m (1×10^13^ vg/mL) and AAV-PHP.eB-CaMKIIα-frChRmine-Kv2.1-oScarlet (5×10^12^ vg/mL) in cold PBS. For mPFC-targeted MA experiments, cortex-wide GCaMP8m expression was achieved by retro-orbital injection (100uL, 1×10^13^ vg/mL), and 300 nL of AAV8-CaMKIIα-ChRmine3.0-Kv2.1-mScarlet (1×10^12^ vg/mL) was delivered locally by stereotaxic injection into bilateral prelimbic cortex (A/P +2 mm, M/L ±0.3 mm, D/V –2 mm from bregma) at a rate of 100 nL·min^-1^ using a 34-gauge beveled needle attached to a 10 µL syringe (World Precision Instruments).

### Surgery

Mice were anaesthetized with 4-5% isoflurane and maintained with 1–2% isoflurane in oxygen and administered sustained-release buprenorphine (0.5mg/kg) subcutaneously for postoperative pain management. The scalp was surgically removed, hydrogen peroxide (3%) was used to remove the periosteum, then the skull was dried before affixing a custom steel head-plate using dental cement (C&B Metabond, Parkell). The skull was then sealed with a thin layer of cyanoacrylate glue (Krazy Glue, Toagosei America Inc.) and clear nail polish (Hard as Nails, Sally Hansen) to render underlying cortex optically accessible. Mice were allowed to recover for at least one week prior to recording or behavioral experiments.

### Widefield Ca^2+^ imaging

Cortical activity was recorded using a custom tandem-lens widefield macroscope comprising Nikkor 50 mm f/1.2 lenses (Nikon), a 495 nm longpass dichroic, and a Kinetix sCMOS camera (Teledyne) operating at 62.5 Hz. A 520/35 nm bandpass filter isolated GCaMP fluorescence. Excitation at 470 nm (Ca^2+^-sensitive) and 405 nm (isosbestic) was strobed on alternating frames for hemodynamic artifact correction. Widefield imaging data were analyzed using custom Python code following established procedures. Data were de-interleaved to separate the signal (470nm) and isosbestic (405nm) color channels, motion corrected, then normalized fluorescence (ΔF/F_0_) was computed for each channel by dividing each pixel’s timeseries by its mean, subtracting 1, then detrending linearly in time. The isosbestic channel was low-pass filtered below 14 Hz and regressed pixelwise onto the signal channel. The resulting residual signal ΔF/F_0_ after regression was warped to Allen Mouse Common Coordinate Framework Reference Atlas version 3 (CCFv3) using an affine transform computed by a least-squares fit between selected keypoints—the center and edges of the olfactory bulb, bregma, and the midline at the most posterior point of RSC—from the individual mouse and the corresponding points on the atlas. For spontaneous activity analysis, pre-processed data were bandpass filtered between 0.5–4 Hz using a zero-phase 3rd-order Butterworth filter. This filtering step was omitted for all-optical and visual detection task analyses, where causal filtering was required to avoid contamination of pre-stimulus periods by post-stimulus activity.

### Convolutional non-negative matrix factorization

Network element discovery used seqNMF, ported to Python with PyTorch GPU acceleration. Processed, atlas-registered, spatially downsampled ΔF/FL movies were first made strictly non-negative by subtracting the minimum value over the entire dataset, then rescaled pixel-wise between zero and one to standardize the range of values. Frames from these movies were then vectorized to produce data matrices 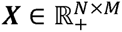 where N = 3225 is the number of pixels and M = 22500 is the number of frames, and then factorized as ***X ≈ W*** ⊛ ***H*** where 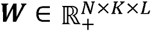 encodes K = 50 spatiotemporal elements of duration L=15 frames (480 ms) and 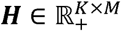 their temporal loadings. The robust regularization framework in seqNMF ensures that redundant factors are eliminated, and each run of the algorithm on a single 12-minute session typically produced around 30 nonzero elements. The value of L was chosen by screening a range of values and taking the shortest duration that produced cohesive bumps of activity that qualitatively matched the data without showing signs of multi-phasic mixed responses.

To generate a set of representative factors for a given session, 100 runs of seqNMF were performed using different random initializations, yielding C candidate elements. Pairwise cross-correlations were computed among all candidate elements across all runs, and the peak cross-correlation value was used as a similarity metric between each pair. From these similarities, a weighted shared nearest neighbor graph was constructed by computing the Jaccard index among the k = 15 nearest neighbors of each candidate element. Louvain community detection^65^ was then applied to this C×C weighted adjacency matrix to identify clusters of similar elements. Within each cluster, the most representative element was identified as the candidate with the highest within-cluster degree z-score^66^. All remaining elements in the cluster were temporally aligned to this representative using the lag of their peak cross-correlation, then averaged to produce a single session-level consensus element per cluster. To combine elements across datasets within a cohort, the graph-based clustering approach was repeated using the set of all session consensus elements as the candidates for the group-level elements. Group level elements were visualized and manually merged in cases of clearly duplicated elements with subtle differences in, e.g., noise signatures. All raw consensus components and merge decisions are presented in (see Extended Data Fig 1). Element discovery was performed individually within Cux2 (n = 63), Sst (n = 77), VIP (n = 70), and CaMKIIα (n = 59) cohorts. In pharmacological intervention datasets, data from the peak drug phase (5-35 min post injection) were segregated into six non-overlapping 5-min segments and these were treated as the lowest level inputs for the element discovery pipeline.

### Defining Sources and Sinks

Source and sink regions of interest (ROIs) were defined separately for each directional element based on their respective spatiotemporal structures. For source identification, an early-versus-late contrast map was computed by taking the pixel-wise maximum activation across the first 3 frames and subtracting the maximum across the last 3 frames of the element. For sink identification, late-phase activity was contrasted against early-phase activity using element-specific frame windows selected by inspection of the spatiotemporal sequence, with the late window ranging from frames 5–11 depending on the element. In both cases the contrast map was averaged with its left-right mirror image to enforce bilateral symmetry, then thresholded at an element-specific value to identify the dominant activated region. Connected components of the thresholded map were identified using *scipy.ndimage.label*, and the largest relevant component was selected. Its intensity-weighted center of mass was computed, a circular ROI of radius 21 pixels placed at this center of mass, and a contralateral ROI defined by reflecting the center across the midline. For the Visual element source, and the Somatosensory and Visual element sinks, automated center-of-mass localization produced ambiguous results and ROI centers were instead placed manually by inspection of the element spatial map overlaid on the Allen CCFv3 parcellation outline. All ROIs were verified by visual inspection against the corresponding directional element spatial footprint.

### Model comparing cell-type-specific directional elements

To assess whether matched and non-matched cell-type fits were statistically equivalent, we performed two complementary paired tests for each cell-type group. Each dataset was fit with each set of cell-type-specific directional elements, a scalar goodness-of-fit as the median pixel-wise correlation between the data and its reconstruction across all brain pixels. The goodness-of-fit was averaged across non-matching fit groups prior to comparison, yielding a single matched and single mismatched value per replicate. The per-replicate normalized delta [(matched – mismatched) / matched] was then used as the unit of analysis.

We first tested for a difference using a two-sided Wilcoxon signed-rank test on the per-replicate deltas. To affirmatively assess equivalence, we additionally performed a two one-sided t-tests (TOST) procedure with equivalence bounds of ±10%, which we defined *a priori* as the threshold of practical equivalence. In TOST, the null hypothesis is that the true mean delta falls outside the equivalence bounds; rejection of this null constitutes positive evidence that the difference is bounded within 10% of the matched fit. All tests were performed separately per cell-type group with no correction for multiple comparisons across groups, as each group constitutes an independent experimental cohort.

To assess spatial equivalence between matched and non-matched cell-type fits, we performed a pixelwise two one-sided t-tests (TOST) procedure across all brain pixels within a predefined brain mask. For each dataset, we computed a normalized per-pixel delta as the difference between the matched fit correlation and the mean correlation across all non-matched fit groups, divided by the matched fit. These per-replicate delta maps were used to construct a pixelwise t-statistic testing whether the mean delta was bounded within ±10% of the matched fit. The TOST p-value at each pixel was taken as the maximum of the two one-sided p-values, such that p < 0.05 constitutes positive evidence of equivalence at that pixel.

To summarize equivalence across the spatial map, we computed the fraction of masked brain pixels for which the TOST null was rejected (p < 0.05). We then tested whether this fraction was greater than the 5% false-positive rate expected under the global null of non-equivalence using a one-sided binomial test (H0: proportion of equivalent pixels ≤ 0.05).

### Orofacial motion energy computation and behavioral state segmentation

Orofacial behavior was monitored continuously during widefield imaging using an infrared camera (Basler ace acA1440-220um) acquiring video at 125 Hz. Frame-by-frame motion energy was computed as the sum of absolute pixel-wise differences between consecutive frames, yielding a total motion timeseries at the native camera frame rate. To align motion energy with the imaging data, consecutive camera frames were grouped into non-overlapping blocks of 4 and summed, reducing the effective frame rate to 31.25 Hz (matching the Ca^2+^ imaging rate). The resulting downsampled motion timeseries was then interpolated to the exact timestamps of the Ca^2+^ imaging frames using linear interpolation (*scipy.interpolate.interp1d*), producing a motion timeseries registered to the brain imaging clock.

For behavioral state segmentation, the motion timeseries was smoothed using a 10-second rolling average. Windows were classified as high-motion (Active) if their average motion energy exceeded the 75th percentile of the per-session distribution, and as low-motion (Quiescent) if it fell below the 25th percentile. Windows falling between these thresholds were excluded from state-specific analyses.

### Orofacial-state-dependent directional element event rates

Directional element temporal loadings were z-scored across the full session and events were detected as peaks exceeding z = 2.5 with a minimum inter-event interval of 5 frames (0.16 s; *scipy.signal.find_peaks*). Per-window event rates (Hz) for each directional element were computed by counting detected events within each 10-second window and dividing by the window duration. Session-level mean event rates were then computed separately for high- and low-motion windows, and compared across behavioral states.

### Statistical analysis of directional element correlates of orofacial behavioral state

Differences in directional element event rates between Active and Quiescent states were assessed using a two-way ANOVA with motion state and directional element identity as factors (including their interaction), implemented via OLS regression (*statsmodels.formula.api.ols*, Type II sums of squares). Post-hoc paired *t*-tests were performed per directional element, comparing session-mean event rates between Active and Quiescent periods across sessions, with multiple comparisons corrected using the Benjamini-Hochberg false discovery rate procedure (*statsmodels.stats.multitest.multipletests*). Effect sizes were computed as Cohen’s *d*, defined as the mean difference in event rate (Active minus Quiescent) divided by the standard deviation of that difference across sessions.

### State-dependent cortical activity maps

To visualize the spatial distribution of cortical activity as a function of behavioral state, pixel-wise RMS ΔF/F_0_ was computed within each rolling 10-second window across all pixels. Windows were assigned to high- or low-motion states as above, and RMS values were averaged separately across high- and low-motion windows within each session, then averaged across sessions to produce group-level spatial maps.

### Pupillometry

Pupil size and position were estimated from high-speed orofacial videos using a DeepLabCut^67^ model trained to localize four keypoints at the top, bottom, nasal, and temporal margins of the pupil. A circle was fit to these four points the circle-fit library in Python (https://pypi.org/project/circle-fit/), and pupil area was defined as the area of the resulting circle. Frames in which any keypoint had a likelihood below 0.6 were flagged, and the resulting pupil area timeseries was linearly interpolated across flagged frames. The interpolated timeseries was bandpass filtered between 0.02 and 8 Hz using a 3^rd^-order zero-phase Butterworth filter. Directional element-triggered average pupil dynamics were computed by z-scoring the filtered pupil area timeseries and averaging across directional element onset times within each session, then averaging across sessions.

Saccades were detected from the frame-wise displacement of the fitted circle’s center. This displacement timeseries was smoothed by taking a 5-frame causal moving average, and frames with smoothed displacement exceeding the 95th percentile were labeled as saccades. Element-triggered average saccade probability was then computed by averaging the binary saccade timeseries across directional element onset times within each session, then across sessions.

### All-optical system and optogenetic stimulation

The all-optical system extended the widefield macroscope with a digital micromirror device (DMD; DLPLCR4500EVM, Texas Instruments; 912 × 1140 mirrors, 7.6 μm pitch) and a 577 nm laser (5 W; Genesis MX577, Coherent) for patterned one-photon optogenetic control. Short-pass filters at 550 nm and 575 nm were added between the GCaMP bandpass filter and the camera to prevent unwanted laser stimulation from reaching the sensor. Finally, the camera was substituted with an OrcaFlash4.0-V2 (Hamamatsu), and 4×4 spatial binning (image size: 512 × 512 pixels; 26 µm × 26 µm pixel size) was used. The frame rate remained 62.5 Hz. ROIs defined in Allen CCFv3 space were warped to native imaging coordinates each session via affine transformation. Stimulation consisted of a 6 ms laser pulse delivered to a 0.6 mm^2^ circular ROI at the network source or sink site. Optogenetic stimulation pulses were temporally interleaved with sCMOS acquisition, being delivered exclusively during inter-frame intervals (i.e., when the camera was not exposing) to minimize optical bleed-through into the imaging signal.

### Pharmacological modulation of directional element dynamics

#### Directional element temporal loading and event detection

To quantify the time-resolved expression of each directional element across pharmacological sessions, consensus spatiotemporal elements derived from the original CaMKIIα cohort were fit to individual session data, yielding per-session temporal loading timeseries 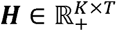 for each of the K = 6 directional elements. Temporal loadings were z-scored relative to a 5-minute pre-injection baseline window by subtracting the baseline mean and dividing by the baseline standard deviation, computed independently for each element and session.

Directional element events were detected as peaks in the baseline-normalized loading timeseries exceeding a z-score threshold of 2.5 and separated by a minimum inter-event interval of 5 frames (0.16 s) using *scipy.signal.find_peaks*. Event times were converted to seconds by dividing by the imaging frame rate and binned into consecutive 1-minute windows. Event rates (Hz) were computed by dividing the per-minute event count by 60.

#### Time-resolved directional element event rates

For group-level analysis, event rate timeseries were computed separately for each session, drug condition (saline, ketamine, diazepam, amphetamine), and directional element. Summary statistics for the "peak drug phase" were computed by averaging event rates across all 1-minute bins falling within the window 5–35 minutes post-injection, then averaging across bins within each session to yield a single session-level summary value per element. This window was chosen to capture the period of stable pharmacological effect following the onset of drug action, excluding the early rise phase.

#### Cortical activity maps during peak drug phase

To generate spatially resolved maps of bulk cortical activity during the peak drug phase, raw ΔF/F_0_ data were bandpass filtered (0.5–4 Hz, zero-phase 3^rd^-order Butterworth), and root-mean-square (RMS) ΔF/F_0_ was computed pixel-wise during the 5–35-minute post-injection period and then averaged across sessions within each drug condition.

#### Statistical analysis of event rates

To test for significant drug-induced changes in directional element event rates, a one-way ANOVA was performed per directional element with drug condition as the independent variable and session-mean peak-drug-phase event rate as the dependent variable, with mouse identity included as an additional factor. Post-hoc pairwise comparisons against saline were conducted using Dunnett’s test (*scipy.stats.dunnett*), which controls the family-wise error rate for multiple comparisons against a single control. Effect sizes were quantified as Cohen’s *d*, computed using the pooled standard deviation across the drug and saline groups. Mouse-level normalization was also performed (subtracting each mouse’s within-saline mean event rate per element) as a sensitivity check; results were consistent across both approaches.

#### Linear Discriminant Analysis decoding of drug identity

To assess whether the multidimensional pattern of directional element event rates was sufficient to identify drug condition, Linear Discriminant Analysis (LDA) classifiers were trained and evaluated on minute-by-minute event rate data. For each 1-minute time bin, session-level event rate vectors were assembled into a feature matrix 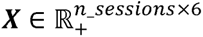, where each of the 6 features corresponded to the event rate of one directional element. Drug classes were balanced by downsampling to the minimum number of sessions across conditions at each time bin. Data were split 67%/33% into training and test sets using stratified random sampling to preserve class balance, with the same random state applied across time bins to maintain comparability of splits over time.

A total of 1000 bootstrap repetitions were performed per time bin. In each repetition, a new balanced subsample was drawn, and a new stratified train/test split was applied. An LDA classifier was fit to the training set and evaluated on the held-out test set. Classifier performance was quantified using the weighted one-vs-rest area under the receiver operating characteristic curve (AUROC; *sklearn.metrics.roc_auc_score*, multi_class=’ovr’, average=’weighted’). A permutation null distribution was generated by repeating the same procedure with shuffled class labels.

A confusion matrix was computed by concatenating true and predicted labels across all repetitions and time bins, normalized by row (i.e., by true label) to express classification accuracy per condition.

#### Cross-time decoding

To assess the temporal stability of pharmacological signatures in the directional element event rate space, a cross-time generalization analysis was performed. For each pair of time bins (m_train, m_test), an LDA classifier was trained on balanced data from minute m_train and tested on separately sampled balanced data from minute m_test, yielding a time × time matrix of AUROC values. Balanced subsamples for training and test were drawn independently. This procedure was repeated for 1000 bootstrap repetitions and results averaged across repetitions.

#### LDA structure coefficients

To characterize which directional elements most strongly drove pharmacological discrimination, a summary LDA was fit on session-level mean event rates from the peak drug phase (5–35 minutes post-injection), pooled across all drug conditions. Features were standardized prior to fitting using z-scoring (*sklearn.preprocessing.StandardScaler* fit on the pooled data). Structure coefficients were computed as the Pearson correlation between each element’s standardized event rate and each linear discriminant axis score across sessions, yielding a matrix of shape (6 elements × 3 LD axes). Unlike raw classifier weights, structure coefficients are scale-free and reflect the contribution of each element to the emergent discriminant axes in interpretable units.

### Visual detection task

Water-restricted mice were trained to lick in response to a faint, isoluminant, drifting Gabor patch (PsychoPy; 1.5 s duration, on a 17.78cm-diagonal screen placed 10 cm from the left eye) to receive a water reward. Stimulus size and contrast were decreased progressively as performance plateaued. Final stimuli had spatial frequency 0.0625 cycles / degree, temporal frequency 2.0 Hz, size 30°, and duration 1.5 s. A 1–3 s no-lick period was enforced before each stimulus onset; off-target licks triggered an aversive air puff. In optogenetic stimulation experiments, network source or control ROIs were optogenetically stimulated for 250 ms (5 pulses at 20 Hz) prior to stimulus onset on pseudorandomly selected trials. Task control used a Bpod r2 finite-state machine (Sanworks) synchronized with the widefield imaging system. Task performance was analyzed using paired t-tests per motif (pre-stimulus analysis of hits vs misses), two-way ANOVA for reaction times, and Generalized Estimating Equation logistic regression for accuracy, with Bonferroni or Holm multiple-comparison corrections as appropriate.

### Projection mapping

Anterograde viral projection data were obtained from the Allen Mouse Brain Connectivity Atlas. Analysis was restricted to injecting the fluorophore into wild-type or Cre into reporter mice (Ai75(RCL-nT)). Injection sites were chosen corresponding to identified motif sources. High-resolution (25 um) projection and injection density volumes were processed using the Mouse Connectivity Cache. To account for variability in viral expression, projection densities were normalized by the total injection signal, defined as the sum of all voxels within the injection site exceeding a **0.05** density threshold. Top-down (dorsal) 2D maximum intensity projections (MIPs) were generated for each experiment. Normalized 2D projections were then averaged across all selected experiments to produce a representative connectivity map. The average injection site was overlaid as a mesh.

Specifically, experiments included are:

MS source: 180601025, 112935169, 180718587, 114292355, 100141495, 112229814, 126852363, 148964212, 112791318 (n=9)

MS sink: 100141454, 141603190, 569993539, 112952510, 657424207, 572753855, 649362978 (n=7)

AL source: 157654817, 180717881, 114290938, 112373124, 112936582, 112162251, 100149969 (n=7)

AL sink: 180709942, 180673746, 180719293, 170721670, 100140756, 157710335 (n=6)

AS source: 127866392, 112951804, 112882565, 126907302, 100142655, 100141473 (n=6)

AS sink: 100141454, 141603190, 569993539, 112514202, 477435412, 112952510, 657424207, 572753855, 649362978 (n=9)

V source: 561918904, 307321674, 307743253, 116903968, 585761377, 309003780, 567723369, 277712166, 571653937, 277714322, 307137980, 277713580, 523180728, 561918178, 589065144, 304564721, 304762965, 563205064, 560965104 (n=19)

V sink: 571652998, 475829896, 146593590, 139426984, 607059419, 573977678, 571401645, 139520203 (n=8)

MA source: 141603190, 528511254, 569932566, 112514202, 475733382, 477435412, 623286726, 112458114, 523481517, 649362978 (n=10)

MA sink: 126861679, 100148503, 112229103, 560045081, 100141599, 591168591, 561986735, 617901499 (n=8)

### Histology and immunohistochemistry

Mice were deeply anesthetized and transcardially perfused with 4% paraformaldehyde (PFA) in PBS. Brains were post-fixed in 4% PFA overnight at 4°C and subsequently sectioned sagittally at a thickness of 50 um using a vibratome. For immunolabeling, free-floating sections were incubated with a rabbit anti-GFP conjugated to Alexa Fluor 488 (Invitrogen; 1:750 dilution) in blocking buffer overnight at 4°C. Sections were counterstained with DAPI and mounted. Representative images were acquired using a confocal microscope to verify fluorophore expression and anatomical localization.

### ASAP7y voltage imaging preprocessing

Raw ASAP7y voltage imaging data were acquired at 133 Hz. Because ASAP7y fluorescence decreases with membrane depolarization, pixel timeseries were sign-inverted prior to all subsequent processing. Data were spatially masked to cortical pixels using a manually defined strict cortical mask registered to the Allen CCFv3. To isolate activity in specific frequency bands, masked pixel timeseries were bandpass filtered using a 6th-order zero-phase Butterworth filter. Four frequency bands were analyzed: delta (0.5–4 Hz), theta (4–10 Hz), beta (10–30 Hz), and gamma (30–60 Hz). For gamma-band analysis, data were additionally spatially downsampled by a factor of 2 (from 8×8 to 16×16 pixel binning) prior to filtering to improve SNR.

For all bands except delta, the analytic envelope of the bandpass-filtered signal was extracted by computing the magnitude of the Hilbert transform (`scipy.signal.hilbert`). The resulting envelope timeseries was then bandpass filtered a second time (0.5–4 Hz, zero-phase, 3rd-order Butterworth) to remove high-frequency fluctuations in the envelope and retain only slow modulations of band-limited power. For delta-band data, no envelope extraction was performed and the bandpass-filtered signal was used directly. Preprocessed data were resampled from 133 Hz to 31.25 Hz using polyphase resampling (`scipy.signal.resample_poly`, up/down ratio 125/532 after simplification by GCD) to match the frame rate of the calcium imaging data.

